# The mechanical nature of injury shapes the composition and temporal dynamics of inflammation after traumatic spinal cord injury

**DOI:** 10.64898/2026.11.18.767599

**Authors:** Chaimae Ahmanna, Alice Gaussot, Kévin Boussion, Kadia Kante, Elias Kanaïe-Atrian, Sylvia Soares, Ysander von Boxberg, Fatiha Nothias

## Abstract

The pathophysiology of traumatic spinal cord injury (SCI) is a complex, evolving process that begins with a mechanical trauma. Progressive secondary damage then occurs, in which neuroinflammation plays a pivotal role. Much has been learned about the persistent inflammatory response during the chronic phase of SCI. However, variations in lesion models, animal species, strains, and the post-lesion intervals investigated make it difficult to conclude on how inflammation contributes to recovery or to exacerbation of damage.

Here, we systematically compared two preclinical rat models of thoracic SCI – contusion and bilateral dorsal hemisection (bDH)– that produce severe injuries leading to comparable initial locomotor deficits. We present a detailed analysis of neuroinflammatory temporal dynamics from acute (1-7 days), subacute (14 days), and chronic stages (> 4 weeks). The two models shared some inflammatory patterns, but specific cellular and molecular responses differed in magnitude, composition, and temporal evolution. These signature inflammatory dynamics were associated with differences in endogenous axonal plasticity, astroglial remodeling and progressive lesion maturation, and with distinct functional recovery rates. As the lesion gradually became chronic, inflammatory activity was remodeled, but persisted. These results reveal the interplay between the mechanical cause of SCI, the corresponding immune response, and the ensuing tissue remodeling processes. They highlight the importance of injury context and subsequent evolution. We discuss them as contributions to our understanding of the physiopathology of spinal cord lesions, with the broader goal of designing therapeutic strategies aiming to appropriately modulate inflammation

**Highlights:**

- Injury mechanics shape inflammation dynamics after SCI
- Inflammatory signatures differ between injury models
- Inflammation dynamics differ despite similar functional recovery
- Axon plasticity mirrors inflammation dynamics
- Chronic inflammation is progressively remodeled

## INTRODUCTION

Traumatic spinal cord injury (SCI) caused by external mechanical impact that compresses, stretches, or transects of the spinal cord tissue, inevitably provokes focal damage to neural cells and fiber tracts. The vasculature is also disrupted, compromising the blood-spinal cord barrier. In addition to these effects, a sustained secondary injury cascade develops, which can be divided chronologically into acute, subacute and chronic phases (Alizadeh et al., 2019). The acute phase is characterized by the initial vascular damage (Figley et al., 2014), combined with an overabundance of extracellular glutamate leading to excitotoxicity (Agrawal & Fehlings, 1997), excess lipid peroxidation (Braughler et al., 1985), and dysregulated ion homeostasis (Teng & Wrathall, 1997; Rosenberg et al., 1999). The subacute phase starts with the activation of apoptotic pathways, leading to demyelination and Wallerian axon degeneration, alongside changes to the extracellular matrix composition. The chronic phase starts once a glial/fibrotic scar forms around the lesion site. It is associated with the progressive stabilization of the lesion, characterized by cystic cavitation, axonal dieback, and further neurodegeneration, with loss of gray and white matter in and around the lesion (Oyinbo, 2011; Badhiwala et al., 2019).

Neuroinflammation plays a pivotal role in the underlying pathophysiology of the evolution of SCI and its long-term outcome, through local and systemic effects. Although inflammation is a physiological process, essential for debris clearance and initiation of the repair processes, excessive infiltration of immune cells can aggravate neural damage (Tran et al., 2018; Hellenbrand et al., 2021). The magnitude of neuroinflammation ultimately depends on the nature, severity, and duration (i.e., compression time) of the primary insult, as well as its evolution (DiSabato et al., 2016; Hellenbrand et al., 2024; Chen et al., 2026)

Inflammation response following SCI involves an interplay between resident and peripheral immune cells. This response has been extensively studied and a certain number of processes have been described. For example, microglia – immune cells resident in the central nervous system (CNS) – rapidly respond to the injury and initially play a protective role (Popovich et al., 2002; Bellver-Landete et al., 2019). They also contribute to pro-inflammatory processes, releasing cytokines that activate astrocytes (Liddelow et al., 2017; Bretheau et al., 2022) and recruit peripheral immune cells, starting with neutrophils and followed by monocyte/macrophages (Popovich et al., 2002; Kubota et al., 2012; Zivkovic et al., 2021).

Together, these cells orchestrate the inflammatory response and influence the balance between tissue damage and repair (Zhang & Gensel, 2014). Monocyte/macrophages share overlapping phenotypic features with activated microglia (Bechmann et al., 2005; Shechter et al., 2009), and, although a strict M1/M2 (Kigerl et al., 2009) dichotomoy is now questioned (Akhmetzyanova et al.2019; von Boxberg et al., 2022), both cell types can be classified as either pro-inflammatory (“M1” cells) or anti-inflammatory (“M2” cells) when activated by alternative pathways. Inflammation is often sustained after SCI, impairing tissue repair (Zhou et al., 2014; Gensel & Zhang, 2015; Bretheau et al., 2022). Indeed, pro-inflammatory activated microglia and macrophages tend to predominate. In response to signals from these cells, lymphocytes – B-cells and various T-cell subtypes – are recruited to the lesion site, with influx typically peaking several days or weeks after the SCI. Complex interplay between the innate and adaptive immune responses, with the secretion of cytokines, chemokines and growth factors that variously amplify and/or resolve inflammation, contributes to both the acute and chronic phases of SCI (Alizadeh et al., 2019). Consequently, the balance between and temporal regulation of these processes critically influence SCI outcomes.

The kinetics of expression and secretion of inflammatory mediators following SCI have been investigated in numerous studies. As a result, we understand some of the factors that lead to excessive or persistent inflammation in the chronic phase (for review see, Bloom et al., 2020; Hellenbrand et al., 2024). This understanding has been extended thanks to recent advances in single-cell and spatial transcriptomic approaches, revealing extensive cellular heterogeneity and distinct molecular states among immune and glial populations after SCI (Li et al, 2022; Wang et al., 2022; Skinnider et al., 2024; Mejia and Horner, 2026). Nevertheless, the temporal organization of inflammatory responses after SCI remains the subject of debate (Hellenbrand et al., 2024; Wrobel et al., 2025). The lack of consensus largely stems from the heterogeneity of experimental approaches, lesion models, timelines, model species, strain, and injury severities (Popovich et al., 1997; Kigerl et al., 2006; Sroga et al., 2003).

The main preclinical models of SCI – contusion and transection – cause fundamentally different patterns of tissue disruption (Siegenthaler et al., 2007). Contusion produces extensive, spatially heterogenous tissue damage (Sharif-Alhoseini et al., 2017; All & Al-Nashash, 2021), whereas transection results in more focused disruption around the lesion site (Dusart & Schwab, 1994; Siegenthaler et al., 2007). These distinct patterns may lead to differences in inflammatory responses, affecting tissue remodeling and endogenous (time-limited) neural plasticity (Soares et al. 2007; Milbreta et al., 2014; Chedly et al., 2017). To understand how different SCI paradigms shape the organization and evolution of neuroinflammation, and to identify potential therapeutic targets (Hellenbrand et al., 2021), we must distinguish the cellular and molecular patterns in each scenario.

In the present study, we systematically compared the inflammatory and tissue responses induced following thoracic SCI. We compared the two main preclinical models of SCI in rats. Inflammation was analyzed during the acute (1-7 days), subacute (14 day), and chronic (28-56 days) post-injury phases. Relationships with transient axonal plasticity, astroglial remodeling, lesion maturation, and spontaneous locomotor functional recovery were assessed. Results show that the mechanical nature of the initial injury shapes the neuroinflammation profile and is associated with differences in tissue remodeling and neural plasticity.

## MATERIAL AND METHODS

### Ethics statement

All surgical procedures and postoperative care were performed in accordance with French legislation and European Union Directive 2010/63/UE. Accordingly, the number of animals was restricted as far as possible, and all efforts were made to minimize discomfort. The study protocol was approved by the Institutional Animal Care and Ethics Committee of the Sorbonne University (#32722-021082012331600 v5).

### Animals and experimental groups

A total of 119 female Wistar rats (Janvier, France) weighing between 230 and 250 g were used for this study. Animals were housed in groups of three under controlled temperature (22 ± 1°C), relative humidity (55 ± 10%) and light-dark cycle (12 h:12 h). Before and after surgical interventions, food and water were available *ad libitum*. Rats were randomly assigned to one of four groups: contusion group, bilateral dorsal hemisection (bDH) group, and control (intact animals) group. For cytokine analysis, an additional group of animals was added, which underwent only laminectomy, without spinal cord damage.

### Spinal cord injury

Surgical procedures were performed at the thoracic level under reversible, continuous Isoflurane® anesthesia (2%-2.5%), and appropriate sterile precautions were used throughout. To perform the SCI, skin and musculature was cut around T7-T10, and the dorsal surface of T8-T9 exposed by laminectomy.

#### Contusive SCI

Contusive SCI was performed using a Benchmark ^TM^ impactor device, applying the approach described in Beliard et al. (Beliard et al., 2022). Briefly, after laminectomy, the spine was stabilized using clamps attached to the base of the impactor device to fix vertebral bodies rostrally and caudally to the impact area. The exposed dorsal surface of the spinal cord was then impacted by a 2.5 mm diameter tip dropped from a height of 7 mm at a speed of 1.96 m/s. The tip was retracted after 100 ms. Following the injury, muscle layers were sutured and the skin was closed.

#### Bilateral Dorsal Hemisection SCI

A bilateral dorsal hemisection (bDH) was performed as described previously (Chedly et al., 2017). Briefly, the spinal cord was exposed by making a longitudinal incision in the dura mater. A hemisection was then made using micro-scissors. To ensure complete ablation of dorsal fiber pathways, an ophthalmic micro-scalpel (blade angle 15°, Micro Feather no. 715) was passed three times through the lesion. Following the injury, the wound was closed as described for the contusion model.

### Post-surgical care

After surgery, rats were housed on extra-soft bedding (Bioservice) to prevent development of friction sores, and their health status was monitored by regular weighing and visual inspection of the surgical wound. The analgesic Buprenorphine (Buprecare® 0.01 mg/kg) was administered before surgery and twice daily for the three days after surgery. To prevent urinary infections, rats were treated daily for the first week post-surgery with subcutaneous injections of antibiotics (Baytril® 10%). Urine was manually evacuated twice daily by abdominal compression until spontaneous micturition was restored. Outcomes were monitored at different days post-injury (dpi): 1 dpi, 3 dpi, 7 dpi, 14 dpi, 28 dpi and 56 dpi.

### Behavioral testing

Locomotor activity was assessed using the standard Basso, Beattie, and Bresnahan (BBB) locomotor scale (Basso et al., 1995) to monitor the immediate impact of lesions and subsequent recovery as well as to assess general well-being. Assessments were performed at 1 dpi and 3 dpi, then weekly until 56 dpi for each group. Animals were placed in an open field study area, where they were carefully visually monitored. Their activity was recorded for 4 min on video. BBB scores were evaluated independently by three trained observers who were blinded to the particular treatment the animals had received. Final scores were calculated as means of the scores assigned by the three observers.

### Time series

For immunohistochemistry and biochemical analysis, at each time point after surgery (1 dpi, 3 dpi, 7 dpi, 14 dpi, 28 dpi and 56 dpi), the spinal cord segment containing the lesion was dissected and prepared. Rats were sacrificed by administering a lethal dose of pentobarbital (Euthasol® 400 mg/mL; 150 mg/kg). They were then transcardially perfused with 0.9% saline at 37 °C, followed by 4% paraformaldehyde (PFA) in 0.1 M phosphate buffer.

### Protein extraction

To allow protein samples to be used for both Western blotting and cytokine array analysis, protein extraction was performed in detergent-free buffer. Rat spinal cord samples (∼1 cm of tissue spanning the lesion site) were homogenized in PBS supplemented with protease inhibitor mix (Sigma; 1/100). Each protein extract was split in two, to be used for Western blotting and Cytokine array testing. At least 3 to 5 rats were used per condition. For 56 dpi (not included in quantitative data), only two specimens were available due to a technical problem. Due to the absence of detergent in the extraction buffer, on Western blots, protein bands occasionally showed some smearing. probably caused by the high level of lipids in CNS tissue.

### Western blotting

Briefly, one aliquot of each homogenate (30 μL) mixed with an equal volume of 2x Laemmli buffer was loaded on 9% or 13% ultra-thin polyacrylamide gels (von Boxberg et al., 2022). After electrophoresis, proteins were transferred to nitrocellulose membranes. Membranes were blocked in a solution of 5% fat-free milk powder in PBS for 1 h before incubating overnight at 4 °C with primary antibodies. After washing, membranes were incubated with alkaline phosphatase-coupled secondary antibodies for 2 h. Binding was revealed by enzymatic reaction using BCIP/NBT reagents (Roth-Sochiel, France).

Primary antibodies used were: goat-anti-MPO (R&D systems, 1/1000), goat-anti-Iba-1 (Wako, 1/1000), rabbit-anti-Arginase-1 (Thermofisher, 1/500), rabbit-anti-CD86 (Thermofisher, 1/1000), mouse-anti-ED1 (Sigma, 1/500), goat-anti-CD206 (Thermofisher, 1/500), and rabbit-anti-glial fibrillary acidic protein (GFAP) (Agilent-Z033401-2, <1/1.000.000). In line with the 3R principle, to reduce the number of animals required, Western blots were usually used to detect two or more antigens. To account for the different reactivity of antibodies, the low levels of certain antigens, and to avoid cross-reactivity, blots were cut into strips covering the approximate molecular weight of the target antigen, as determined using a pre-stained MW marker (Thermo Scientific™ PageRuler™ Plus, 10-250 kDa). The intensity of specific bands was quantified using ImageStudioLite software (LiCor). To normalize for protein loading, intensities were expressed relative to staining with mouse-anti-GAPDH (Santa Cruz, 1/1000) or mouse-anti-actin (Santa Cruz, 1/500).

### Cytokine array

Aliquots of homogenate were centrifuged after thawing. Cytokine and chemokine expression was measured using Rat Cytokine Array Panel A (R&D Systems, Minneapolis, MN, USA) according to the manufacturer’s instructions. Briefly, nitrocellulose membranes spotted with the cytokine/chemokine array were blocked for 1 hour before incubating with spinal cord homogenates containing 600 µg protein (BCA assay, Biorad) were added and incubated overnight at 4 °C. Membranes were then washed before incubating with the biotin-labeled detection antibody cocktail for 1 hour. After washing, binding was revealed by incubating with streptavidin-HRP for 30 minutes, washing once again and applying a chemiluminescent agent (provided with kit Panel A; R&D systems, Minneapolis, MN). Relative cytokine levels were determined using Fiji/ImageJ software.

### Immunohistochemistry

For immunohistochemistry, spinal cord segments were post-fixed overnight in the PFA perfusion solution, and cryopreserved in sucrose (15% for 24 hours, then 30% for 48 hours) at 4 °C. After embedding in OCT (Tissue-Tek), a series of horizontal sections – thickness 40 μm, for imaging, 20 μm for cell counting – were cut using a Leica CM3050 S cryostat. Sections were mounted on slides and stored at-80 °C. For immunostaining, tissues were permeabilized with 0.3% Triton X-100 in PBS for 5 minutes at room temperature, and blocked for 1 hour in 10% bovine serum albumin (BSA) / PBS before exposing to primary antibody diluted in 5% BSA/PBS overnight at 4 °C. Primary antibodies used were: Rabbit-anti-GFAP (Agilent-Z033401-2, 1/2000), mouse-anti-GFAP (Sigma-G3893, 1/2000), mouse-anti-β3 tubulin (Tuj-1) (Biolegend - 801202, 1/2000), mouse-anti-ED1 (CD68; Sigma, 1/100); rabbit-anti-CD206 (M2-macrophage marker; Santa Cruz, 1/500); rabbit-anti-CD86 (M1-macrophage marker, Thermofisher - MA532078, 1/200), rabbit-anti-Myeloperoxidase (MPO) (Thermofisher-PA5-16672, 1/200), mouse-anti-CD4 (BioRad-MCA153GA,1/500), mouse-anti-CD8, (BioRad-MCA48GA 1/500), rabbit-anti-IBA-1 (Fujifilm - 019-19741, 1/200). Nuclei were counterstained with DAPI (Euromedex, 3617980).

Images of large horizontal spinal cord sections were acquired on a Zeiss Axio Zoom V16 microscope and combined using image stitching. High-magnification images were acquired on a Zeiss Confocal LSM 980. All images were processed using ImageJ (NIH, USA) and Adobe Photoshop CS6.

### ED1 and Iba1 immunostaining: quantitative analysis through ROI segmentation

The quantification strategy was inspired by the method described for mouse SCI (Brennan et al., 2022). It was adapted, optimized, and partially automated for use with rat spinal cord. Images from horizontal spinal cord sections double-stained for Iba1 and ED1 were acquired at a final magnification of 100× on a Zeiss Axio Zoom V16 microscope equipped with image stitching software. Identical exposure times were set for all samples when acquiring images for the same marker. Quantification was based on up to four sections for each animal, depending on tissue availability and staining quality; 3 to 5 animals were used for each time point in each SCI model. Original microscope images were processed using a semi-automated multi-step pipeline (illustrated and described in full in **supplementary figure 1**).

### CD4+ and CD8+ T-cell counts

For both SCI models, at 7, 14, and 28 dpi, CD4⁺ and CD8⁺ cells were counted on four 20 μm thick spinal cord sections per animal (n = 3 per group). Cells were manually counted in five predefined areas of interest (AOIs) within each section: one rostral, one caudal, two lateral, and one central location within the lesion. Only cells showing clear membrane staining and DAPI-positive nuclei were counted. A region of interest (ROI) corresponding to the lesion was defined based on the GFAP-negative area. Each AOI within this ROI was a square representing 6% of the total ROI. Together, the five AOIs covered 30% of the lesion area. The average number of cells across the five AOIs was then multiplied by 16.67 (100/6) to estimate the total number of cells within the entire ROI. For each section, estimated cell numbers were normalized to a reference area of 0.2 mm². The average of the normalized values obtained from the four sections was then plotted for comparison.

### Axon density measurement

For each animal, four sections at the level of the injury site (spaced 160 µm apart) were double-stained for GFAP and b3-tubulin at specific post-injury time points. Using Fiji/ImageJ software, an ROI was delineated based on the GFAP-negative area, that also corresponds to the area of injury devoid of neuron cell bodies. Invasion of this area by axons was measured based on b3-tubulin staining intensity. The mean value per rat was calculated, and compared between SCI models at 7 dpi, 14 dpi, and 28 dpi.

### Statistical analysis

Statistical analysis was performed and graphs were produced using GraphPad Prism software. Due to limited sample sizes, data normality was assumed but could not always be formally tested. Two-way ANOVA was used for all quantifications. When the interaction between time and lesion type was significant, post hoc multiple comparisons (Sidak’s or Tukey’s test) were used to assess group differences at each time point and/or to assess temporal changes within each group. When no significant interaction was detected but a main effect of time or group was present, appropriate post hoc tests were used to assess changes over time (combining all groups), or differences between groups (combining all time points). Data are presented as mean and standard deviation (SD). Statistical details for each dataset are reported in the respective figure legends.

## RESULTS

### Spontaneous functional recovery: validation of lesion severity

Lesion severity was assessed and compared between cohorts by analyzing spontaneous functional recovery based on results from the open field test and the corresponding BBB score. As shown in **supplementary figure 2**, in both models, rats had initially completely lost hindlimb motor function. Recovery was more extensive and faster after bDH than after contusion injury, but both groups reached a plateau at approximately 28 dpi.

### Immune response

In parallel to these functional assays, we focused our attention on the immune response within the lesion, and compared the effects of the two lesion types on cellular and molecular profiles. The immune response relies on a complex interplay between early responders – innate immune cells – and later involvement of myeloid cells, closely followed by lymphoid cells. Chemokine and cytokine signaling contribute to regulating these processes.

### Early innate immune response: neutrophil recruitment patterns in SCI models

Neutrophils are key players in the early innate immune response, and once activated express myeloperoxidase (MPO). In both SCI models, activated neutrophils accumulated in the damaged spinal cord rapidly after injury (**Fig.1 A**). Western blot analysis of relative amounts of MPO (**Fig.1 B**) confirmed that activated neutrophils peaked at 3 dpi in both SCI models, with higher levels in contusion than bDH lesions (p=0.0565). Thus, between 3 and 7 dpi, MPO levels dropped significantly (p < 0.0001 for both groups), and were barely detectable at 14 dpi (**Fig.2**).

**Figure 1:**
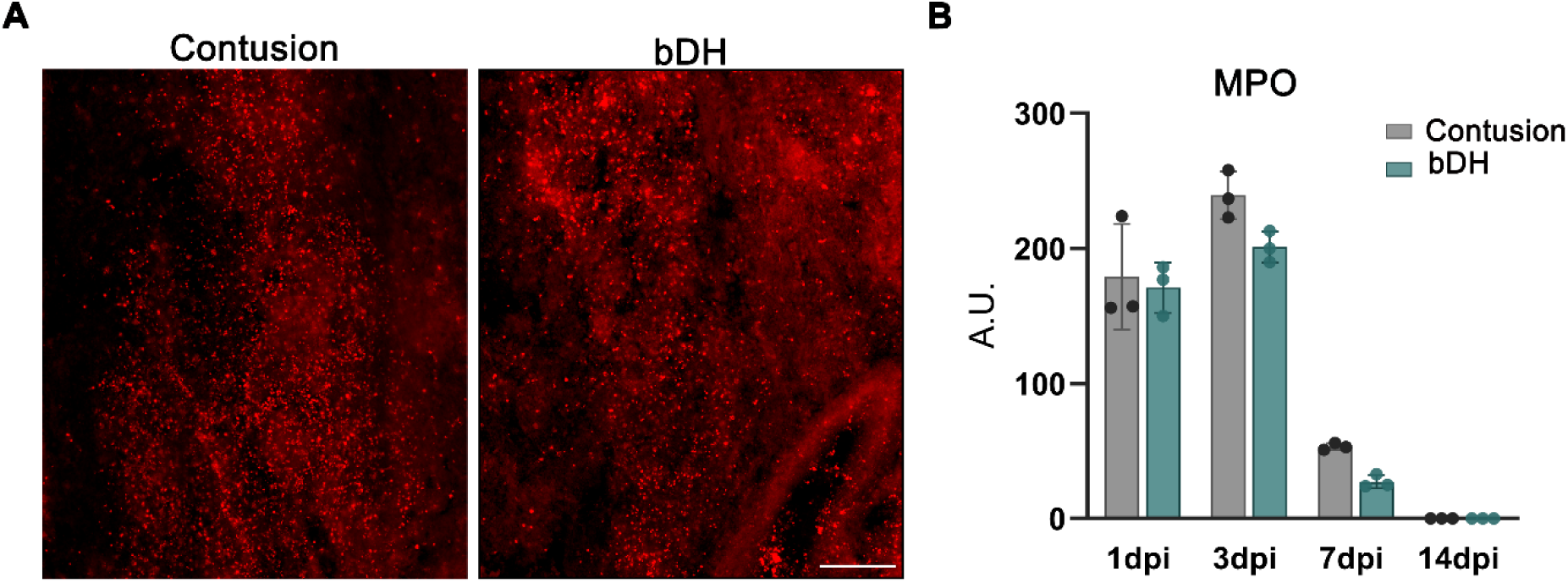
Temporal dynamics of neutrophil marker protein myeloperoxidase after SCI. A: Immunohistochemical labeling of spinal cord sections corresponding to the lesion center following contusion or bDH injury. Sections from 3 dpi were labeled for myeloperoxidase (MPO-red) (scale bar =500 µm). **B:** Relative MPO expression at 1, 3, 7, and 14 dpi, as assessed by Western blotting (a representative Western blot of spinal cord protein extracts labeled for MPO can be viewed in Fig. 2). Two-way ANOVA was used to analyze effect of time (p < 0.0001) and lesion type (p = 0.0198) on MPO expression. Tukey’s post hoc test was performed to assess differences across time points (pooled across groups), revealed a significant decrease in MPO expression between 3 and 7 dpi (p < 0.0001). As only two lesion types were compared, the significant main effect of lesion type reflects a global difference between these two groups across all time points. Results are expressed as mean, error bars correspond to SD (3 animals per condition). bDH: bilateral dorsal hemisection. A.U: Arbitrary Unit, dpi: days post-injury.

**Figure 2:**
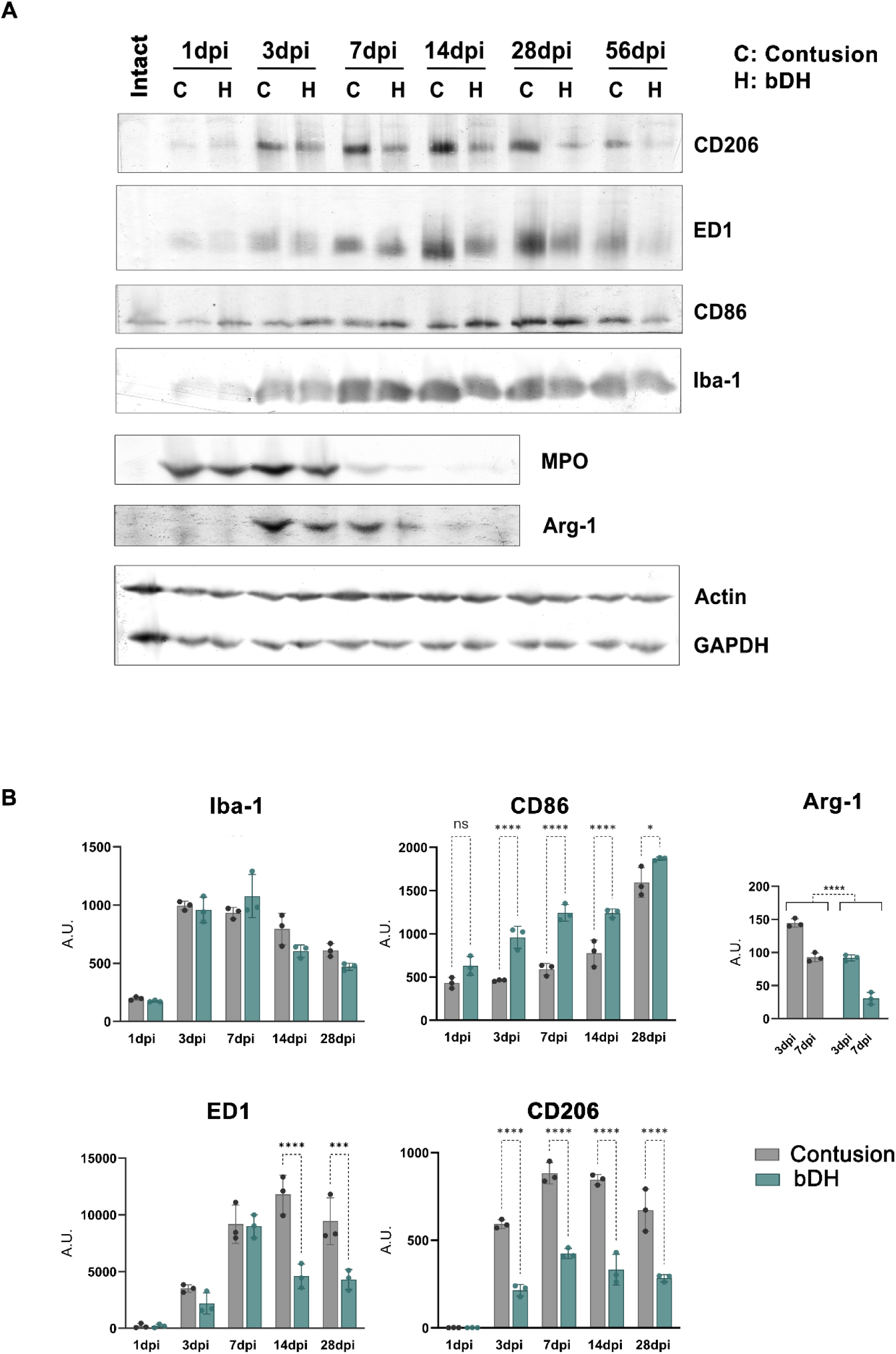
***Differential expression of microglia and macrophage marker proteins in two models of SCI* A**: Representative Western blot of spinal cord protein extracts labeled for CD206, ED1, CD86, Iba-1, MPO, and Arg-1. Actin or GAPDH staining levels were used for signal normalization. Samples were taken from intact spinal cord and at 1, 3, 7, 14, 28, and 56 dpi following contusion or bDH injury. **B:** Quantification of Western blot signals for macrophage and microglia marker protein expression. Results are expressed as mean, error bars correspond to SD (3 animals per condition). Two-way ANOVA was performed for all quantifications. Multiple comparisons were performed between lesion types at each time point, and over time within each group. Post hoc analyses were restricted to within-group temporal comparisons. *p*-values from Sidak’s post hoc tests are indicated on the graphs where appropriate. Statistical significance: (* p< 0.05, ** p< 0.01, **** p< 0.0001). Note that quantitative analysis of MPO, a neutrophil marker, is included in Fig.1; in intact SC, Iba-1 was not detected under the immunoblot conditions used in this study.

### Myeloid immune response: microglia/macrophage recruitment

Microglia, CNS-resident myeloid immune cells, share several morphological and functional features with periphery-derived macrophages. Both cell types express the ionized calcium-binding adapter molecule 1 (Iba-1) (Ohsawa et al., 2000), (Imai et al., 1996; Carlson et al., 1998; Sasaki et al., 2001), and, following activation, the lysosomal protein ED1 (Kullberg et al., 2001).

In intact spinal cord, Iba-1-positive, ED1-negative microglia with a ramified phenotype – long processes and small cell bodies – were distributed throughout white and gray matter (**Fig.3**). Following SCI, before 3 dpi these cells remained largely negative for ED1 and mainly exhibited a ramified morphology (data not shown). Iba-1 immunostaining increased at 7 dpi, and density of Iba-1+ / ED1+ cells increased strongly at the lesion epicenter (**Fig. 3**). These cells acquired a more rounded (less ramified) morphology, whereas Iba-1+ / ED1- cells located outside the lesion epicenter retained a stellate, ramified morphology (**Fig. 3**).

**Figure 3:**
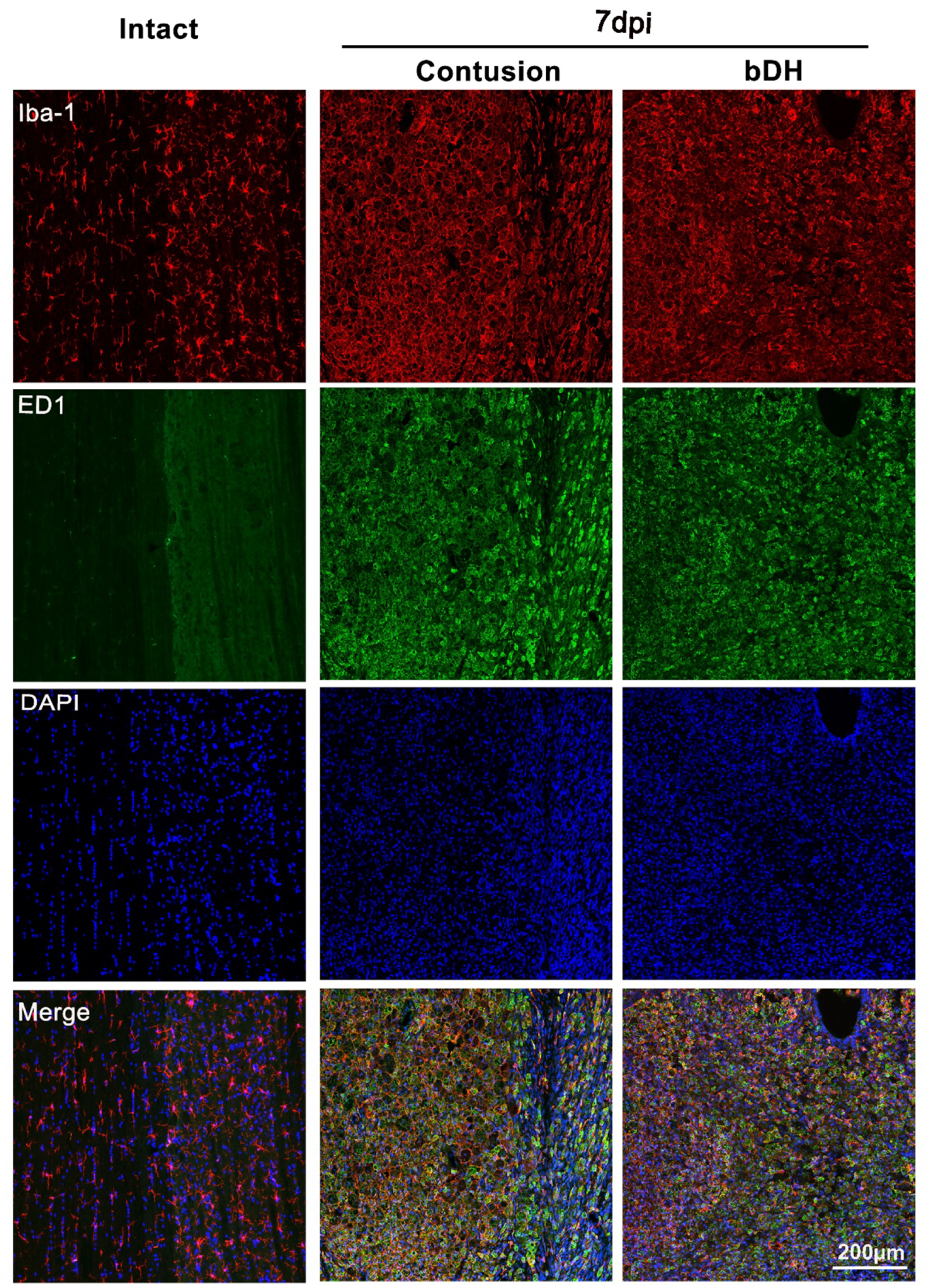
Microglia/macrophage morphological changes and phagocytic macrophage response in contusion and bDH SCI models. Representative confocal images showing microglia activation and macrophage invasion into the lesion epicenter. Sections at 7 dpi were double labeled for Iba-1 (red) and ED1 (green) in contusion and bDH SCI models, and in intact spinal cord tissue. Nuclei were counterstained with DAPI (blue).

Western blot analysis revealed further details on the temporal profiles of Iba-1 and ED1 expression (**Fig. 2A-B**), and differences between the two SCI models. Iba-1 was detectable from 1 dpi, and increased markedly thereafter, reaching maximal levels between 3 dpi and 7 dpi (**Fig.2 A, B**; comparing levels at 1 dpi and 3 dpi, p < 0.0001) in both lesion models. Although levels remained high up to 56 dpi (not quantified) compared to 1 dpi, protein levels progressively decreased after 7 dpi. This decline appeared more pronounced in the bDH model, with a significant reduction between 7 dpi and 14 dpi (p < 0.0001). ED1 protein levels showed a similar trend (**Fig.2 A, C**). ED1 was barely detectable at 1 dpi in either lesion model, but increased thereafter, reaching a peak at 7 dpi for the bDH model and 14 dpi for the contusion model. Levels remained elevated in the contusion model up to 28 dpi, but decreased rapidly in the bDH model. The difference between the two SCI models was significant at 14 dpi (p <0.0001) and 28 dpi (p<0.0002).

In the bDH model, the more rapid decline in Iba-1 and ED1 protein levels mirrored redistribution of ED1+ cells, which were gradually restricted mainly around the lesion epicenter and along median fiber tracts (**Fig. 4**). In contusion lesions, ED1+ cells remained more widely distributed. At 28 dpi, despite the formation of cystic cavities in both lesion models (**Fig. 4A**), Iba-1+/ED1+ cells were still present within the lesion core. The density of ED1+ cells appeared higher after contusion (supplementary figure 3), in line with Western blot results showing sustained ED1 expression (**Fig. 2**). High-magnification images revealed Iba-1+ cells in contusion lesions to have an amoeboid morphology (**Fig. 4A**), whereas ramified cells were visible around bDH lesions.

**Figure 4.**
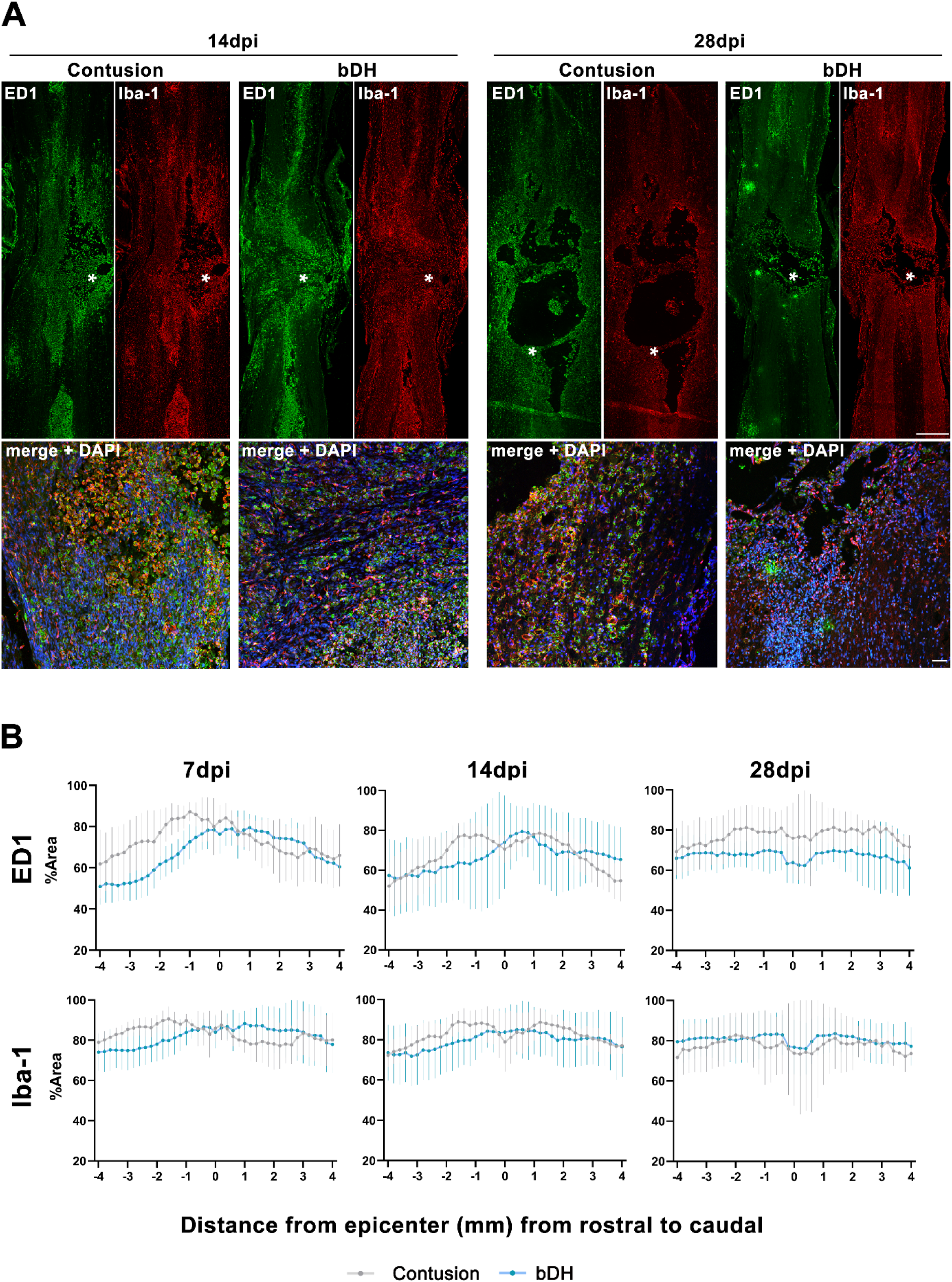
Spatiotemporal distribution of microglia and macrophages at the lesion site in contusion and bDH SCI models. A: Representative immunohistochemistry images of spinal cord sections from contusion and bDH models at 14 dpi (left) and 28 dpi (right) labeled for Iba-1 (red), ED1 (green), and DAPI for cell nuclei (blue in merge photo). Each panel displays both whole-section views (top, scale bar: 500 µm; rostral, upper, caudal, lower part) and higher magnification confocal images (bottom, scale bar: 100 µm). Asterisks indicate the locations where high-magnification images were acquired (lesion center at 14 dpi and lesion cavity border at 28 dpi). **B:** Quantitative analysis of immunostaining distribution of ED1 and Iba-1 cells through ROI segmentation of spinal cord from lesion epicenter (axis, 0) to adjacent segments, rostral (axis, 0 to-4 mm) and caudal (axis, 0 to +4) at 7, 14, and 28 dpi. Two-way ANOVA or mixed-effects analysis was performed for all quantifications. Sidak’s post hoc comparisons were applied to compare results between groups. For the contusion model, data were obtained from four animals at 7 dpi, five at 14 dpi, and three at 28 dpi. For the bDH model, five animals were analyzed at 7 dpi, four at 14 dpi, and five at 28 dpi.

### Myeloid immune response: inflammatory profiles

We next examined whether the specific lesion type influences the inflammatory phenotype of these microglia/macrophage populations. The pro-inflammatory marker CD86 was detectable from 1 dpi, and progressively increased up to at least four weeks post-injury in both models. However, levels were significantly higher in the bDH model than in the contusion model (3 dpi p<0.0001, 7 dpi p<0.0001, 14 dpi p<0.0001, and 28 dpi p=0.0154; **Fig. 2A-B**). At 56 dpi, although not quantified, CD86 levels appeared to decline in both lesion models but remained higher compared to 1 dpi.

The anti-inflammatory markers Arginase-1 (Arg-1) and CD206 displayed different temporal profiles. In both models, Arg-1 was barely detectable at 1 dpi, it rapidly increased at 3 dpi before dropping once again from 7 dpi (**Fig. 2A**). Although the time-course was similar in both lesion models, Arg-1 expression was significantly higher in the contusion model at 3 dpi and 7 dpi (p < 0.0001; **Fig 2**). CD206 was detectable from 3 dpi, peaking at 7 dpi in both lesion models. Like Arg-1 levels, CD206 levels were significantly higher in the contusion model, at all time points from 3 to 28 dpi (p < 0.0001; **Fig.2B**). CD206 protein expression appeared to be maintained for longer in the contusion model; it was barely detectable at 56 dpi in the bDH model (**Fig.2A**). Immunohistochemistry corroborates these differential polarization patterns, clearly showing the changes in CD86 and CD206 expression levels at 7 dpi and 28 dpi (**Fig. 5**).

**Figure 5:**
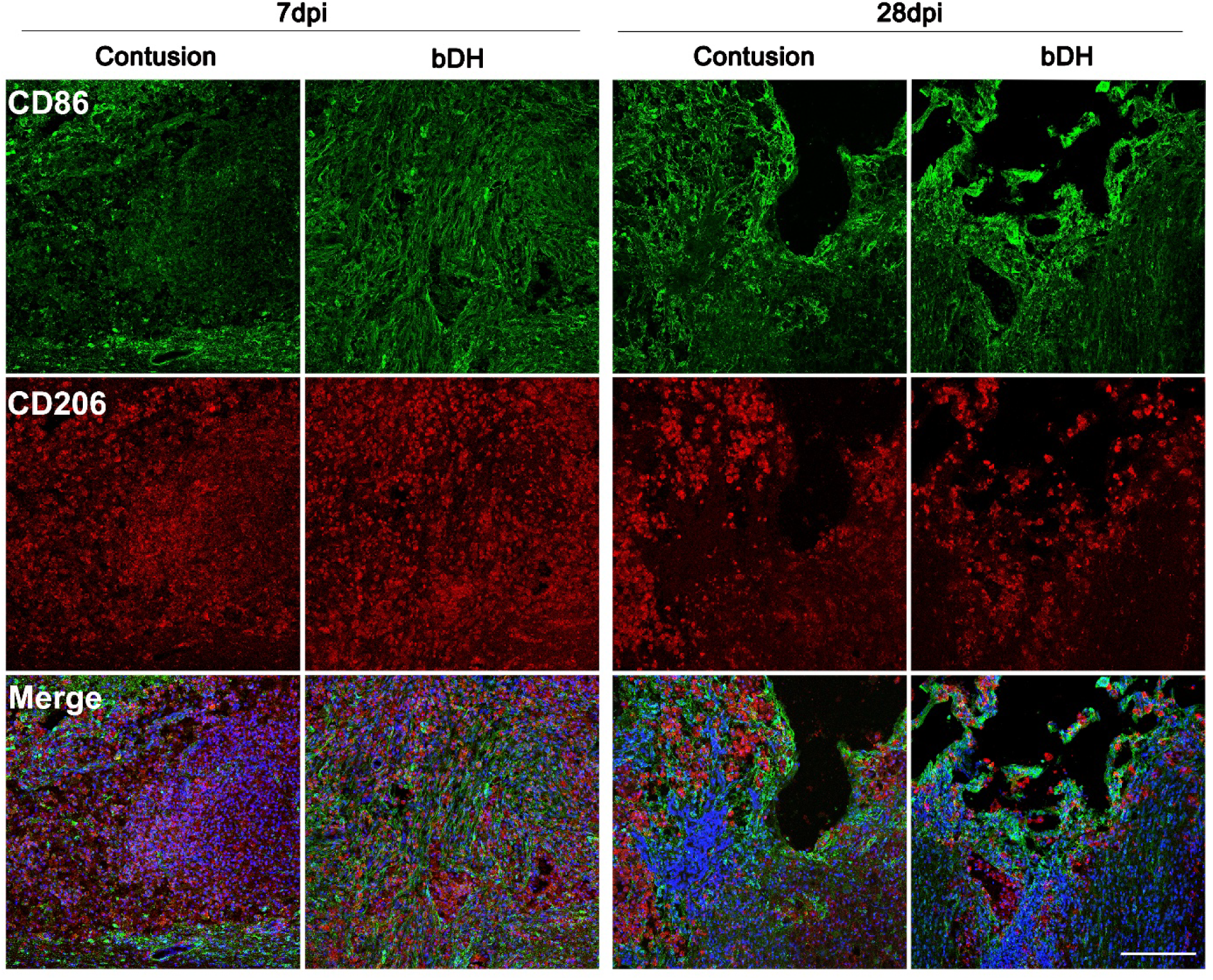
Distribution of polarized macrophages at the lesion site after contusion and bDH SCI at 7 and 28 dpi. Representative immunohistochemical images of sections stained for CD86, M1 pro-inflammatory macrophages (green) and CD206, M2 anti-inflammatory macrophages (red) at 7 dpi at the lesion site, and around the lesion cavity at 28 dpi, with DAPI staining (blue in merge images) for cell nuclei. Scale bar: 200 µm.

The lesion type influenced not only the duration of microglia/macrophage activation, but also their inflammatory phenotype. bDH lesions were associated with a stronger and more sustained pro-inflammatory response, whereas contusion injuries had a more pronounced and longer-lasting anti-inflammatory response.

### Adaptive immunity in the injured spinal cord: T-cell infiltration profiles

We next investigated the adaptative immune response by analyzing the distribution of CD4+ and CD8+ T lymphocytes within the lesion site at different time points. T-cell distribution was heterogeneous in both bDH and contusion SCIs, where it closely followed the shape and the extent of the lesion (**Fig. 6A**). Although T-cells were also detected outside the GFAP-negative area delineating the lesion, our analysis exclusively focused on the cells within the GFAP-negative area. This analysis revealed some differences between the two SCI models (**Fig. 6B**).

**Figure 6:**
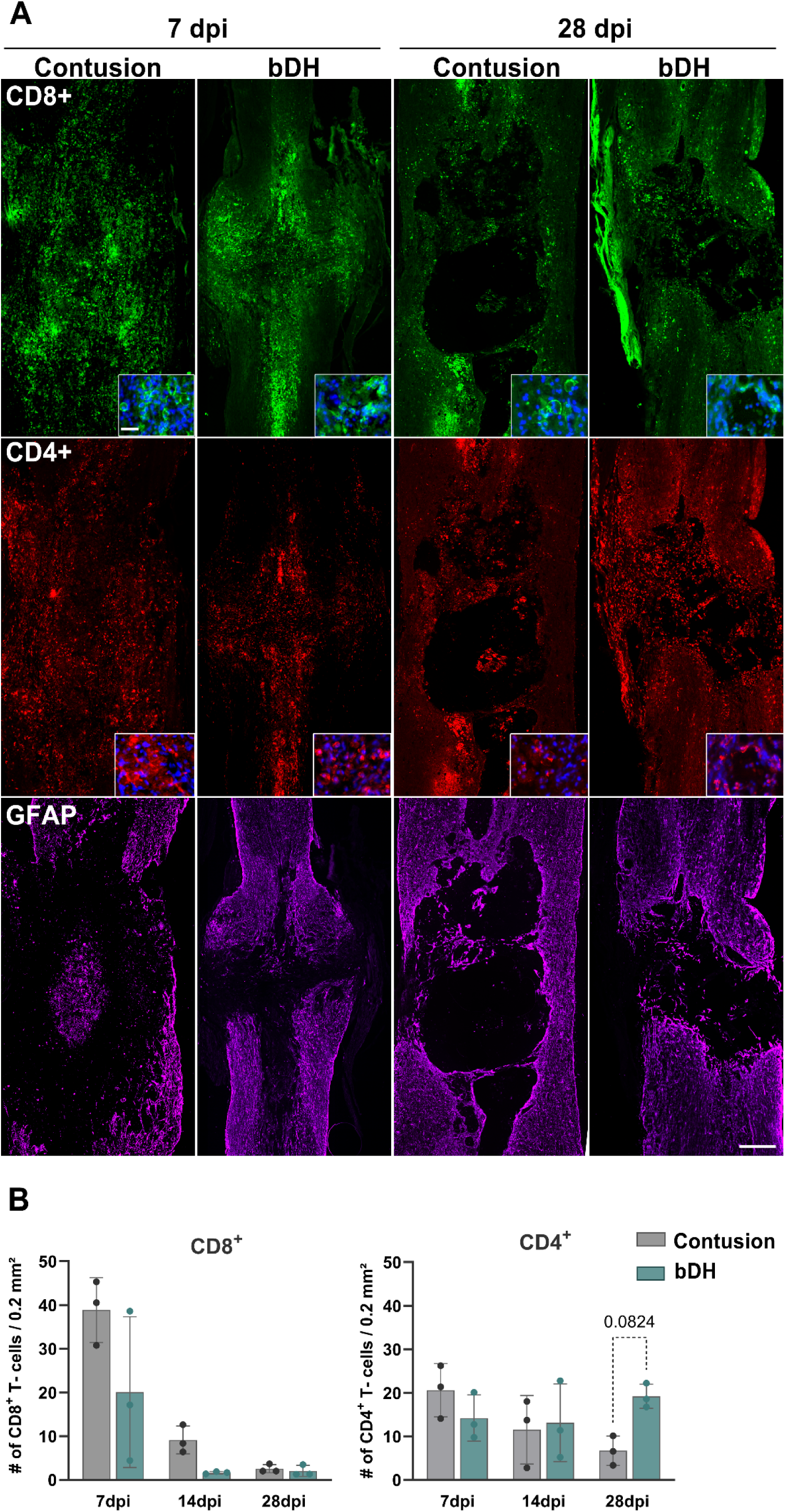
Lymphocyte infiltration pattern in the two SCI models. Immunofluorescence staining and corresponding quantitative analysis of CD4+ and CD8+ T-cells within the lesion site, defined here as GFAP-negative area. **A**: Immunofluorescence staining of spinal cord sections labeled for CD4+ T-cells (red), CD8+ T-cells (green), and GFAP (violet), at 7 dpi and 28 dpi. Scale bar: 500 µm. Insets: high-magnification views of CD4+ / DAPI and CD8+ / DAPI; scale bar: 150 µm. **B**: Quantitative analysis of CD4+ and CD8+ T-cell numbers. Bar graphs represent the number of CD8+ (left graph) and CD4+ (right graph) T-cells per 0.2 mm² of spinal cord tissue within the GFAP-negative area at 7, 14, and 28 dpi. Results are expressed as means, error bars correspond to SD, calculated from 3 animals per condition, and four sections per animal. Two-way ANOVA was performed for both quantifications. For CD8+ T-cells, a significant main effect of time (*p* = 0.0001) and lesion type (*p* = 0.0319) was observed. Tukey’s post hoc test (across time points, both groups combined) revealed a significant decrease in CD8⁺ cell number between 7 and 14 dpi (*p* = 0.0005). For CD4+ T-cells, a trend toward a significant interaction was detected (*p* = 0.0608), which justified performing Sidak’s multiple comparisons test the difference was not significant at 28 dpi (p = 0.0824).

CD8^+^ T-cells were mainly detectable from 7 dpi, which corresponded to their peak. This overall profile was the same in both injury models, although absolute numbers of CD8+ cells were significantly higher in the contusion model (p=0.0319).

CD4^+^ T-cells were also detectable from 7 dpi. Their numbers were similar in the two SCI models at 7 dpi and 14 dpi. However, by 28 dpi, their numbers showed a tendency for decline in the contusion lesion, but not in the bDH lesion (p=0.0501 compared to 14 dpi), = and levels also tended to be higher in bDH compared to contusion at 28 dpi (p= 0.0824).

### Chemokine and cytokine profiles mirror the temporal dynamics of immune cell recruitment

As indicated above, immune cell recruitment and activation is controlled by cytokines and chemokines in the inflammatory environment. We therefore analyzed chemokine and cytokine levels for a broad spectrum of inflammation-related mediators known to participate in SCI pathophysiology (Cizkova et al., 2018; Chio et al., 2019; B. Zhang et al., 202; Hong et al., 2018). The same spinal cord homogenates were used for cytokine arrays and Western blots, by splitting each sample (see methods). Results are categorized based on the reported properties of the chemokines and cytokines (pro-inflammatory, anti-inflammatory, growth factors, inhibitors, adhesion molecules, etc.). Trends and changes over time are shown in graphs (**Fig. 7, 8, and 9**). Levels of molecules were compared not only between the two SCI models, but also between SCI models and intact and laminectomy-only groups (comparison of the two control groups can be found in **supplementary figure 4**). The inflammation fluctuations observed after laminectomy alone confirm the need for this type of control in all studies involving models of SCI. To account for the effects of post-surgery recovery, only statistically significant differences compared to the laminectomy-only group are presented here.

**Figure 7:**
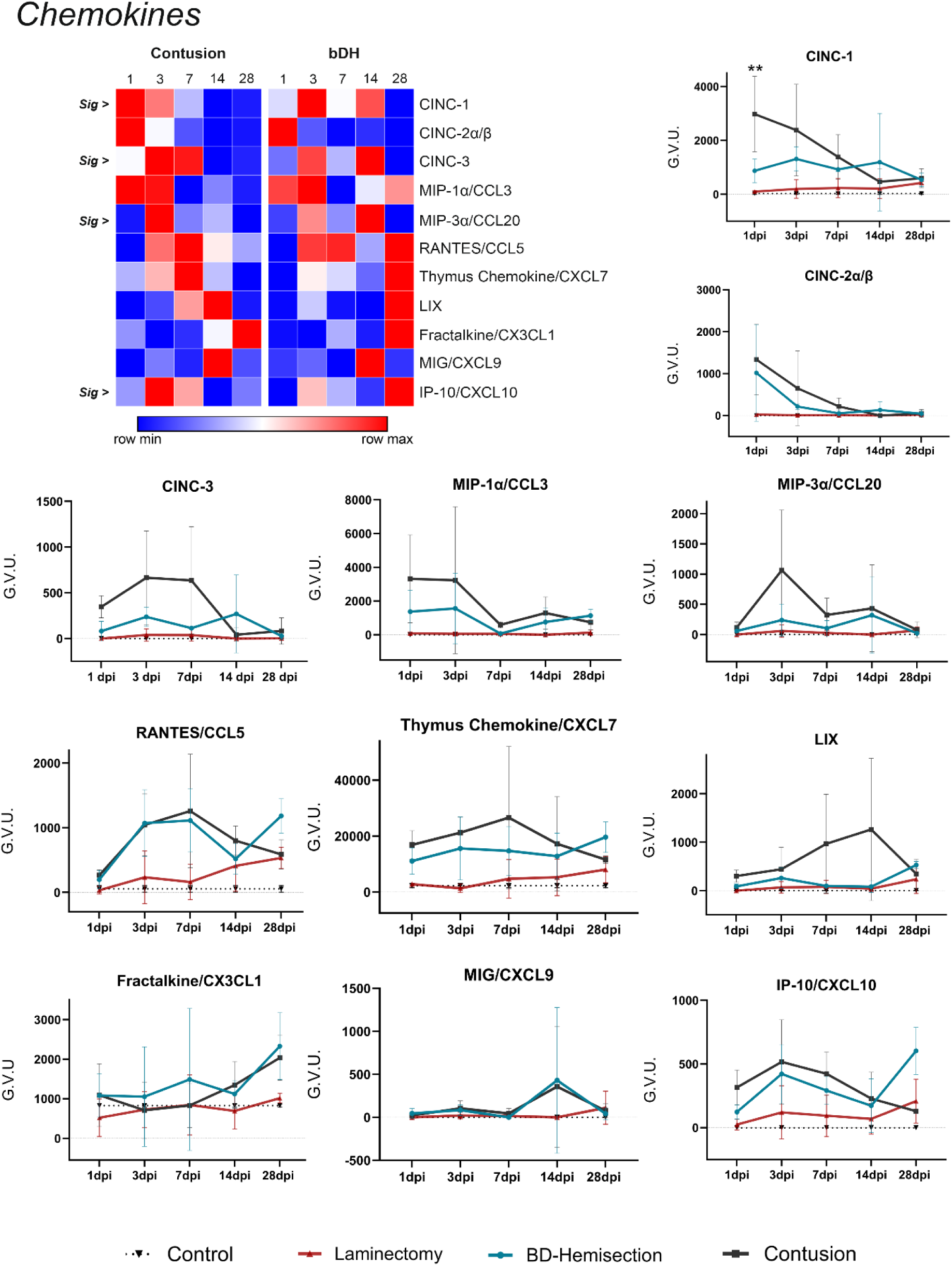
Comparative ELISA analysis of chemokine levels in spinal cord tissue at different time points post-lesion. Top left: Heatmap showing the relative expression levels of various chemokines, based on raw gray value units, with color coding indicating relative expression (blue = minimum (row min), red = maximum (row max)). Graphs show cytokine expression levels in both SCI models compared to laminectomy and control (intact) groups. Values are presented as mean, error bars correspond to SD (n=3/4). Note that graphs show only significant differences between contusion and bDH groups. Additional statistical differences between laminectomy and each SCI model are detailed below. Effects of time, lesion type, and their interaction were assessed by two-way ANOVA; only relevant post hoc comparisons are reported. Number of animals per condition: 3-4.

**Figure 8:**
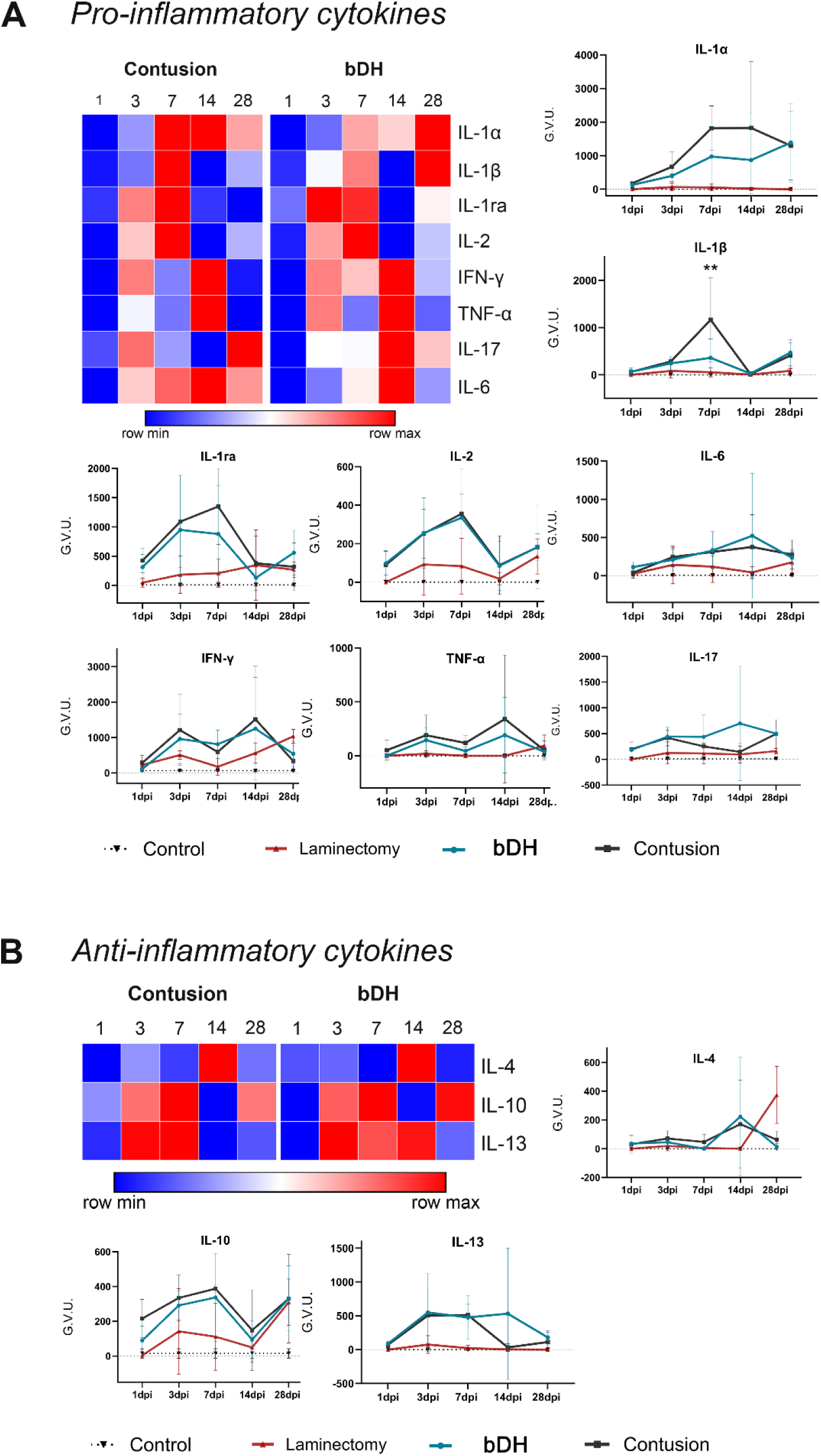
Comparative analysis of pro-and anti-inflammatory cytokine levels in contusion and bDH SCI models across different time points. **A**. **Pro-inflammatory cytokines:** Heatmap illustrates the relative protein levels of pro-inflammatory cytokines in contusion and bDH models. Corresponding graphs show cytokine expression over time in each model, compared to laminectomy-only and controls. **B. Anti-inflammatory cytokines:** Heatmap shows relative protein levels of anti-inflammatory factors IL-4, IL-10, and IL-13, with corresponding time-course graphs for each cytokine in the contusion and bDH models compared to laminectomy-only and controls. Values represent means, error bars correspond to SD (n=3/4). Note that graphs only show data for significant differences between contusion and bDH. Effects of time, lesion type, and their interaction were assessed by two-way ANOVA; only relevant post hoc comparisons are reported.

### Chemokine expression after SCI

### Early chemokine induction parallels early recruitment of innate immune cells

Consistent with the rapid infiltration of MPO-positive neutrophils observed during the acute phase (**Fig. 1**), several chemokines involved in leukocyte recruitment display a robust increase rapidly after SCI (**Fig.7**). This response profile was common to both lesion models, but differences in magnitude and timing were observed.

Thus, Cytokine-Induced Neutrophil Chemoattractants (CINCs), which promote neutrophil infiltration (Q. Li et al., 2021), exhibited the strongest early response. Three members of this family were assayed. CINC-1 levels were significantly higher after contusion than after bDH injury at 1 dpi (p=0.0059; **Fig.7**). Levels subsequently decreased steeply in the contusion model, with the result that the lower level CINC-1 expression was maintained for longer in the bDH model. CINC-2 showed a similar expression pattern in both lesion models. CINC-3 also had a similar profile, with a slight delay, peaking between 3 and 7 dpi in the contusion model, followed by a marked decline by 14 dpi. In the bDH model, overall expression was modest, with distinct peaks at 3 dpi and 14 dpi. Comparison combining all time points revealed higher levels of CINC-2 a/ß (p=0.0371) and CINC-3 (p=0.0025) after contusion compared to laminectomy-only. The differences were not significant for the bDH model.

Other chemokines also showed a prominent early response. Macrophage Inflammatory Proteins MIP-1α (CCL3) and MIP-3α (CCL20) were significantly higher following contusion than following laminectomy-only (p=0.0126 and p=0.0344, respectively; **Fig. 7**). Both chemokines are involved in the recruitment and migration of immune cells, including monocytes/macrophages and various T-cell subsets (Menten et al., 2002; Schutyser et al., 2003; Nandi et al., 2016). In both lesion models, other chemokines involved in inflammation, RANTES (Regulated upon Activation Normal T-Cell Expressed and Secreted, or CCL55), Thymus chemokine (CXCL7), and IP-10 (interferon gamma-induced protein 10 or CXCL10), were also elevated compared to the laminectomy-only control (**Fig.7**).

A later response was observed for LIX (Lipopolysaccharide-Induced CXC chemokine or CXCL5), which peaked at 14 dpi in the contusion model and 28 dpi in the bDH model (**Fig.7**). All times combined analysis revealed a significant difference between the two models(p=0.0498). This chemokine recruits both myeloid and lymphoid cells.

Fractalkine/CXC3L1 was detectable in both control spinal cords, in line with its constitutive expression under physiological conditions. Although levels increased progressively following SCI in both models, peaking at 28 dpi, the differences were not statistically significant (compared with laminectomy, p=0.0826 for bdH and p=0.3138 for contusion).

### Cytokine expression after SCI

#### Pro-inflammatory cytokines

At 1 dpi, levels of pro-inflammatory cytokines were relatively low in both SCI models. Levels then increased progressively, following largely similar profiles: IL-1α, IL-1β, IL-1ra, and IL2 peaked at 7 dpi; IL6 peaked at 14 dpi; both IFN-γ and TNF-α showed a biphasic pattern, with elevated levels at 3 and 14 dpi (**Fig.8A**). Nevertheless, absolute levels were higher after contusion for several cytokines, particularly IL-1β (p=0.002) at 7 dpi. IL-17 was the only cytokine for which the pattern of expression differed between the two models. Overall, IL-17 levels were higher in bDH model, with significant group effect (p=0.0336), and significantly higher levels than in the laminectomy model when all time points were considered (p=0.0256; **Fig. 8A**).

#### Anti-inflammatory cytokines

From 1 dpi, anti-inflammatory cytokines also exhibited a dynamic temporal profile in both models (**Fig. 8B**). Thus, IL-10 and IL-13 levels rose between 3 dpi and 7 dpi, whereas IL-4 is transiently expressed around 14 dpi in the two SCI models. In the laminectomy-only control, IL-4 expression displayed a distinct patterns, behing detectable only at 28 dpi and reaching higher levels than in either of the two lesion models. The increase in IL-13 was sustained for longer in the bDH model than in the contusion model, with a significant difference relative to the laminectomy-only control (p=0.0286). IL-10 was the only anti-inflammatory cytokine to show a similar expression profile between SCI models and laminectomy-only control (**Fig.8 B**).

### Inhibitor, adhesion molecule, and growth factor expression following SCI

Overall, the levels of adhesion molecules, L-selectin, TIMP-1, and sICAM-1, were elevated after SCI in both models (**Fig. 9A**). L-selectin and TIMP-1 declined relatively rapidly, whereas sICAM-1 was sustained until later stages (28 dpi). The response of CNTF, VEGF and GM-CSF growth factors tended to increase during the subacute phase, followed by a general steep decline at 28 dpi in both SCI models. This profile is clearly visible in the heatmaps (**Fig.9B**).

**Figure 9:**
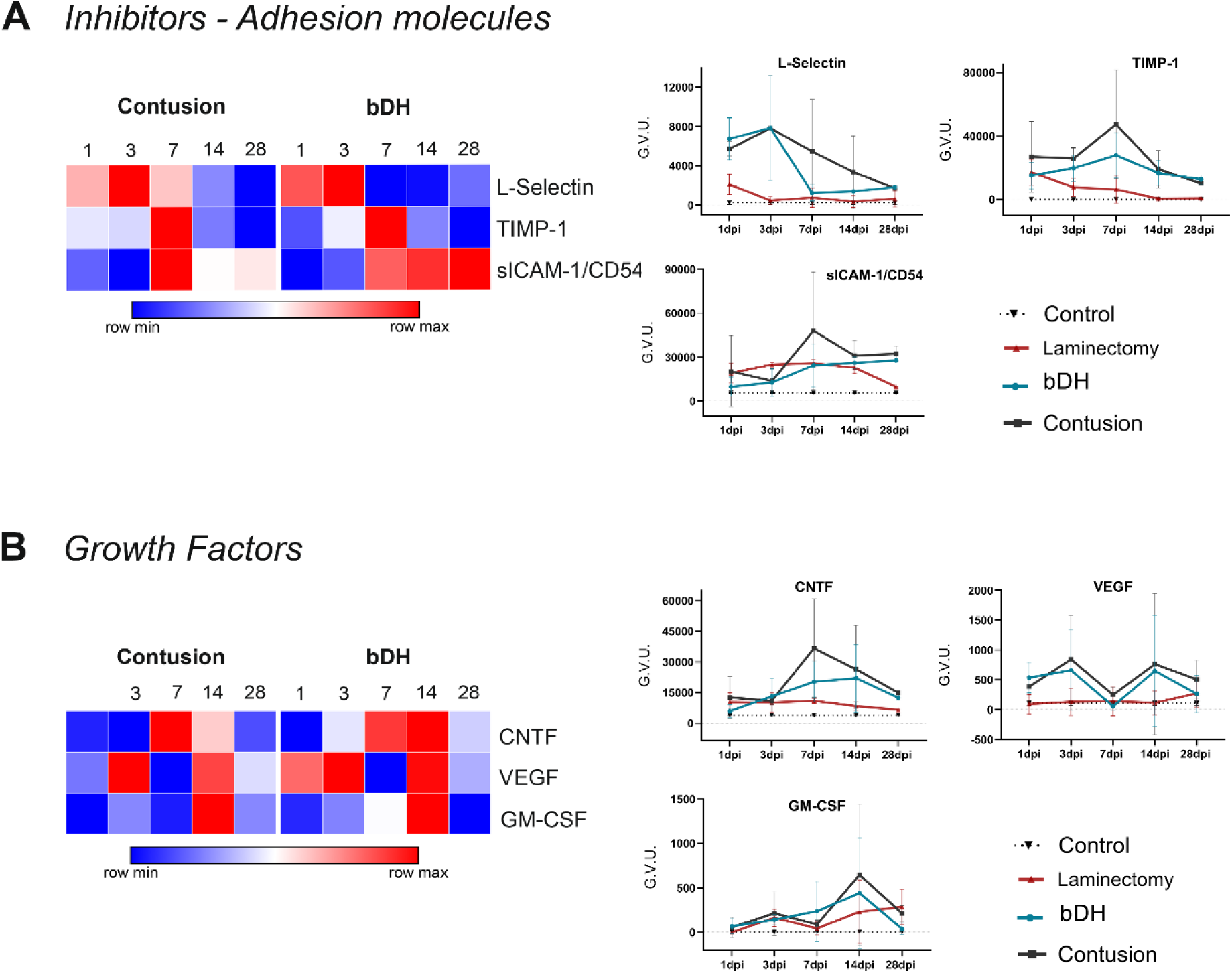
Comparative analysis of adhesion molecules, growth factors, and TIMP-1 in contusion and bDH SCI models across different time points. A. TIMP-1 and adhesion molecules: Heatmap illustrates the relative expression levels of TIMP-1, L-selectin, sICAM-1 in contusion and bDH models. Graphs show the temporal patterns compared to laminectomy and control. **B. Growth factors:** Heatmap shows relative levels of CNTF, VEGF, GM-CSF, with corresponding time-course graphs for each factor in contusion and bDH models compared to laminectomy and control. Values represent mean, error bars correspond to SD (n=3/4). Note that only significant differences between contusion and bDH are shown on graphs. Two-way ANOVA was used to test the effects of time, lesion type, and their interaction; only relevant post hoc comparisons are reported.

### Axonal plasticity and astrocyte remodeling corroborate inflammation dynamics

Inflammatory changes in the context of SCI are of interest in how they relate to tissue remodeling. We examined this aspect of recovery by analyzing axonal plasticity and astrocyte reactivity within lesions.

β3-tubulin (Tuj-1) immunostaining was used to evaluate the proportion of the GFAP-negative lesion area occupied by axons (**Fig.10**). Spinal cord sections taken at 7, 14 and 28 dpi were analyzed. Neuron cell bodies were detected in spinal segments adjacent to the lesion, but the GFAP-negative area was devoid of neurons due to neurodegeneration. At 1 and 3 dpi, axonal density could not be quantified due to the amount of debris related to the primary insult. Between 7 dpi and 14 dpi, axonal density within the lesion area peaked for both models. By 28 dpi it had dropped significantly (contusion: p=0.0001, bDH: p=0.0001). Axons occupied a larger area in the bDH model compared to the contusion model, with the difference reaching significance at 14 dpi (p = 0.0329; **Fig. 10B**).

**Figure 10:**
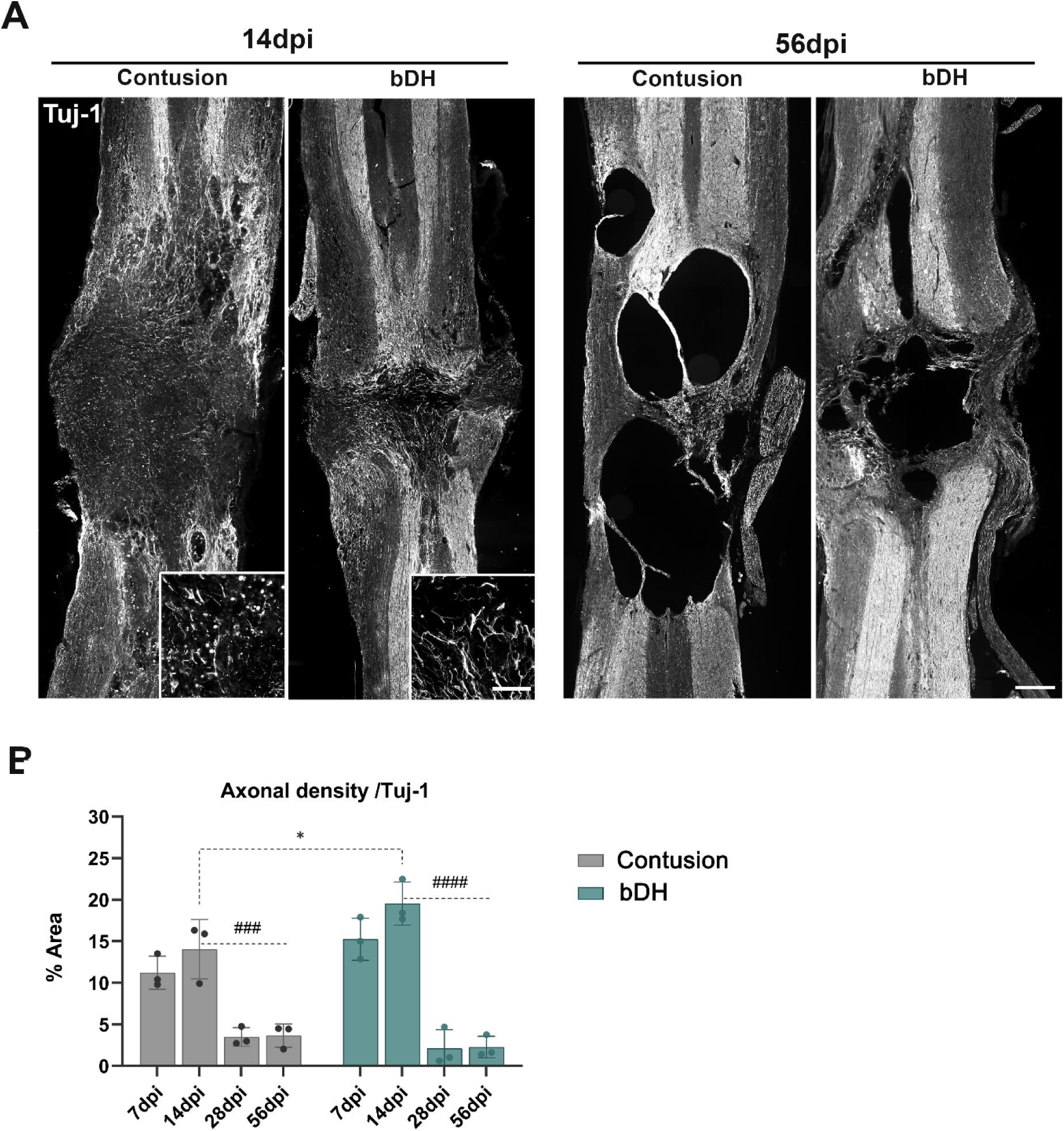
Lesion-induced axon plasticity in response to contusion vs. bDH SCI models. **A**. Axonal labeling of sections from contusion and bDH models. Sections at 14 and 56 dpi were labeled with Tuj-1 (anti-β3-tubulin). Inset: high-magnification view (scale bar: 145 µm). **B.** Relative density of Tuj-1 immunostained fibers within the lesion, expressed as a percentage of the GFAP-negative lesion area occupied by axons. (###, p = 0.0001; ####, p < 0.0001). Statistical analysis was performed using two-way ANOVA revealed a significant main effect of time (*p* < 0.0001) and a significant interaction between time and group (*p* = 0.0290). Sidak’s post hoc test identified a difference between groups at 14 dpi, while Tukey’s post hoc test revealed differences across time points. Values represent means, error bars correspond to SD (n=3-4).

At 56 dpi, the latest time point analyzed, strong loss of axons and a large cystic cavity was observed in both the contusion and the bDH models (**Fig.10A**). This tissue remodeling was accompanied by the maturation of a dense astroglial scar surrounding the lesion (**Fig.6**). Astrocyte reactivity, as measured based on GFAP-protein amounts from Western blots, reached its highest level at 56 dpi in both SCI models, with higher overall levels in the bDH model than in the contusion model (p=0.0001).

## DISCUSSION

Inflammation is a pivotal driver of secondary damage in SCI. Indeed, as it evolves through distinct stages – from acute to chronic – the underlying changes often prevent inflammation resolution (Braughler et al., 1985; Teng & Wrathall, 1997; Rosenberg et al., 1999; Oyinbo, 2011; Badhiwala et al., 2019; Alizadeh et al., 2019). Here, we characterized the complex and dynamic inflammatory response in two well-established preclinical models of SCI: bilateral dorsal hemisection transection (bDH), representing a transection-type injury, and contusion injury. Our results indicate that the ultimate outcome of SCI is not determined by injury severity alone.

The two SCI models were comparable – the lesion was applied to the same spinal segment and had functional locomotor outcomes – and produced similarly severe initial functional deficits (BBB scores). While both models exhibited inflammatory patterns commonly described in the literature, our comparative analysis of the acute, subacute, and chronic phases revealed how these patterns evolved, and highlighted model-specific differences. Our data also highlight an intricate interplay between the characteristics of the initial injury (inflammation dynamics), of the immune response (lesion remodeling), and of the structural adaptation occurring after SCI (neural plasticity).

### The mechanical insult causing SCI shapes the onset and evolution of neuroinflammation processes

The parallels between the two models provide a functional basis for comparing the inflammatory response triggered in each case, while minimizing differences related to the severity of the locomotor deficits in acute post-injury period. BBB scores were used to validate the lesion and to assess spontaneous recovery. Although the maximum BBB score achieved in the bDH model was higher than that in the contusion model, both models reached their spontaneous recovery plateau within a similar time frame, at around 4 weeks post-injury. Thus, the extent of spontaneous functional recovery differed between models, but the time between injury and stabilization was similar. This similarity suggests that injury severity is not the only factor contributing to differences in neuroinflammatory responses between the two SCI models.

Contusion injuries provoke a widespread response due to sustained mechanical stress and tissue compression, including damage to vascular structures (Beliard et al., 2022). Conversely, transection injuries produce strong, but more localized, mechanical disruption, with more limited rostro-caudal spread of secondary damage. These differences in the initial mechanical insult were directly associated with critical events in the early inflammatory response.

In accordance with previous studies (Albadawi et al., 2017; J. Wang, 2018), neutrophils were the first peripheral immune cells observed to infiltrate the lesion. Activated neutrophil presence peaked at 3 dpi and declined thereafter, as previously reported (Taoka et al., 1998; Carlson et al., 1998; Chatzipanteli et al., 2000). In the context of SCI, neutrophils have both beneficial and detrimental effects. Their beneficial impact relies on their phagocytic activity, contributing to debris clearance. However, excessive or prolonged neutrophil activation leads to oxidative stress, protease release and amplification of secondary inflammatory damage (Zivkovic et al., 2021; F. Zhang et al., 2024). The overall neutrophil response observed was more pronounced after contusion compared to bDH. The early neutrophil response was accompanied by marked increase in secretion of neutrophil-associated chemokines, such as CINCs (e.g., CINC-1-3) and the adhesion molecule L-selectin, known to contribute to neutrophil recruitment (Omari et al., 2005; Merabova et al., 2012).

In addition to their own response, neutrophils participate in the subsequent recruitment of peripheral monocytes and macrophages. During the subacute phase of SCI, the early increase in MPO levels and MPO+ cells, correlated with subsequent increases in ED1 and Iba-1 levels – markers of active microglia and infiltrating peripheral macrophages. Between 7 and 14 dpi, cells positive for these two markers were observed. Like neutrophils, macrophage accumulation was more pronounced after contusion and was accompanied by higher associated chemokine levels (MIP-1α/CCL3, RANTES/CCL5, and IP-10/CXCL10). These chemokines signal to T lymphocytes; their infiltration patterns were also similar, peaking around 7 dpi, as reported elsewhere (Wu et al., 2017).

Based on these cellular events, contusion appears to elicit a more robust early immune response than transection injury.

### Different SCI models have distinct cellular and molecular inflammatory signatures

The stronger immune cell recruitment observed after contusion could seem obvious considering the extent of the immediate damage caused. However, closer analysis of the cellular and molecular inflammatory responses revealed a more complex picture. Although peripheral immune cells accumulated in higher numbers after contusion, expression of repair-associated macrophage markers, including arginase-1 and CD206, was also higher during the acute and subacute phases. In the bDH model, a stronger inflammatory macrophage-associated profile was observed, characterized by higher CD86 protein levels and higher numbers of CD86-positive macrophage/microglia at the lesion site.

One possible explanation for the stronger peripheral immune cell recruitment after contusion injury relates to the involvement of meningeal interfaces. The meninges represent an active immunological niche, and act as a key interface between the CNS and the peripheral immune system, controlling immune trafficking and lymphatic drainage (John 2018; Asepelund et al. 2025; Louveau et al., 2018; Song et al., 2020; Gonuguntla and Herz, 2023). Although bDH directly disrupts the meninges at the lesion site, contusion injury produces broader spinal cord deformation that may also affect surrounding meningeal compartments. Vascular alterations following contusion may also contribute to the differences observed between the two models. Indeed, chronic contusion is associated with a preferential reduction in spinal blood volume within the dorsal part of the lesion (Biliard et al., 2022), due to the dorsal-to-ventral nature of the mechanical impact. Thus, in addition to the extent of parenchymal damage, the differential peripheral immune recruitment and distinct immune polarization between the two models may be influenced by the anatomical territories affected by the initial injury.

The outcome of inflammation after SCI is not simply linked to immune cell recruitment. The inflammatory signature, which shapes how resident and recruited peripheral immune cells interact with their environment, must be considered. Reports of distinct neuroimmune responses following different SCI paradigms have been recently published (Chen et al., 2026; Helleenbrand et al., 2024). However, direct comparison is difficult when a single post-injury time point is analyzed (Chen et al., 2026), or when studies only cover a short period not extending to lesion maturation (Helleenbrand et al., 2024).

By analyzing the two lesion models from the acute phase onwards up to the chronic phase, we observed that the differences in organization of the immune response between models were not restricted to myeloid cells. The T lymphocyte response also displayed distinct temporal profiles, with a more pronounced but transient accumulation of CD8+ T-cells after contusion. CD4+ T-cell numbers, in contrast, were more sustained and higher at later stages following bDH lesion. However, like macrophages, CD4+ and CD8+ T-cell populations should not be considered homogenous functional pools. Thus, while the distinct CD4+ and CD8+ T-cell profiles observed here indicate that the type of injury influences the broad class of T-cells recruited and how recruitment evolves, they do not define the functional state or contribution of the recruited T-cells. Indeed, advances in high-dimensional cellular profiling, including multiparametric flow cytometry, single-cell and single-nucleus RNA sequencing, and spatial transcriptomics have revealed substantial heterogeneity in terms of transcriptional states and functional properties (Cao et al., 2022; Chelyshev and Ermolin, 2023). In line with this heterogeneity, the functional contribution of T-cells after SCI appears to be highly context-dependent, with differences in activation states, kinetics of recruitment, and antigen specificity (Moalem et al., 1999; Kipnis et al., 2002; Daglas et al., 2019; Xu et al., 2021). Early studies with T-cell-deficient animals suggested that the adaptative immune response was detrimental to SCI outcomes (Potas et al., 2006). However, this conclusion should be nuanced in light of more recent work demonstrating that specific injury-associated T-cell populations can exert a neuroprotective effect (Gao et al., 2024). Thus, engineered T-cells expressing injury-associated receptors promote neuroprotection, in particular through IFN-γ signaling modulating the local myeloid response.

Depending on the model, the immune response after SCI differs in terms of cellular and molecular composition, temporal evolution, and effects on tissue remodeling processes depending on the initial injury. These findings further support the hypothesis that injury mechanics influence the composition and evolution of the immune response, rather than simply determining its magnitude.

### Inflammatory dynamics are associated with endogenous axonal plasticity, astroglial remodeling, and lesion maturation after SCI

We also observed distinct patterns of tissue remodeling between models. In both lesion models, axon density increased during the early post-injury period, reaching a maximum between 7 and 14 dpi, but had markedly decreased by 28 dpi. This evolution is consistent with the well-established transient endogenous axonal plasticity within the lesion site site and adjacent spinal segments reported in rats following injury (Soares et al., 2007; Milbreat et al. 2014). As the lesion evolves, the extracellular environment becomes less permissive to axonal growth, through the accumulation of inhibitory cues, including CSPG and myelin-associated proteins such as Nogo-A, MAG, and OMgp, and progressive reactive astroglial remodeling (Fitch and Silver, 1997; Milbreta et al., 2014; Schwab ME and SM Strittmatter, 2014; Filous and Schwab, 2018). By comparing the temporal profile of axonal plasticity with that of the cellular and molecular responses to inflammation, we identified a number of temporal parallels.

In both lesion models, the concurrent decrease in L-selectin and in neutrophil numbers coincided with the onset of the active axonal plasticity period, beginning at 3dpi. This observation further supports a potential link between the decrease (resolution) of the early neutrophil response and the transition toward a more permissive environment for active axonal plasticity and subsequent functional recovery from 7dpi. Indeed, downregulation of L-selectin is reported to directly reduce neutrophil accumulation and oxidative stress, improving long-term neurological recovery and white matter preservation (McCreedy et al., 2018). We observed distinct temporal profiles between the two injury models. Thus, L-selectin declined more gradually following contusion, whereas the decrease was more abrupt after bDH. Similarly, LIX/CXCL5, which contributes to neutrophil trafficking (Sepuru et al., 2014), reached significantly higher levels following contusion compared to bDH. The less prolonged LIX/CXCL5 response and the more rapid decline in L-selectin after bDH may have created a more favorable environment for tissue remodeling, and reduced the spread of inflammation into adjacent spinal segments. Given the greater axonal plasticity observed after bDH, we hypothesize that differential regulation of neutrophil recuitment and trafficking could contribute to the difference in the extent of axonal plasticity and functional recovery observed between the two injury models.

The anti-inflammatory cytokines IL-4 and IL-13 have been associated with tissue repair and neuroprotective responses after SCI (David and Kroner, 2011; Gordon and Martinez, 2010, Lima et al., 2017). Here, the highest levels of IL-4 and IL-13 in both lesion models were detected during the period of greatest structural remodeling (7-14 dpi), determined based on axon density within the lesion area. In the bDH model, axonal density was greater at 14 dpi compared to the contusion model, in line with more sustained IL-13 expression and more abundant CD4+ T-cell presence. In accordance with this result, adoptive transfer of CD4+ T-cells after SCI was reported to increase IL-4 signaling, promoting neuroprotection, axonal regrowth, and functional recovery (Amo-Aparicio et al., 2021; Francos-Quijorna et al., 2016; Walsh et al., 2015).

Finally, axonal remodeling coincided with dynamic changes in trophic and tissue remodeling factors. CNTF and TIMP-1 increased during the early-subacute period, and declined at later stages. CNTF has neuroprotective and neurotrophic properties and can be produced by astrocytes and Schwann cells within a spinal lesion (Friedman et al., 1992; X. Liu et al., 2007; Ramaswamy & Kordower, 2012). TIMP-1 is involved in MMP-dependent extracellular matrix remodeling and can modulate neurite outgrowth and morphology, at least in part through regulation of MMP2 activity (Ould-Yahoui et al., 2009; Ries, 2014). The temporally regulated expression of MMP following SCI, together with their association with axonal plasticity, is consistent with active changes in the tissue remodeling during the early and subacute phases following SCI (Goussev et al., 2003).

The above molecular changes were mirrored by alteration in cellular activity. Thus, astroglial reactivity progressively increased, with GFAP expression continually rising at least up to 56 dpi, and reaching higher overall levels after bDH. Similarly, CD86 protein levels and numbers of CD86+ inflammatory macrophages were highest in the late post-injury stage in the bDH (transection) lesion model. The two effects may be linked as factors released by both macrophages and microglia promote astrocyte proliferation and influence the expression profile of reactive astrocytes (Haan et al., 2015; Bellver-Landete et al. 2019), and microglia secrete factors contributing directly to astrocyte proliferation and glial scar formation after SCI (Bellver-Landete et al., 2019).

Evidence for this potential interplay is provided by levels of the chemokine RANTES/CCL5, expressed at high levels by reactive astrocytes following injury (Li et al, 2015). RANTES/CCL5 was increased in bDH lesions at the latest time point analyzed (Fig. 7).

The delayed astroglial response, occurring after the decline in axonal density, is consistent with a progressive maturation of the lesion, leading to cystic cavitation and establishment of a defined glial scar (Soares et al., 2007, Milbreta et al., 2014; Chedly et al., 2017). Our results further show that this structural transition coincides with the plateau of spontaneous recovery in both SCI models at 28 dpi.

Injury mechanics influence the cytokine expression profile and cellular activity within an SCI lesion, leading to distinct functional outcomes upon stabilization, which occurs within the same timeframe regardless of the model.

### Persistent but remodeled inflammatory activity characterizes chronic lesions

Following lesion maturation, cellular and molecular inflammatory activity persisted in both injury models. The neutrophil-associated response largely declined, whereas activated microglia/macrophages remained present alongside CD4+ T-cells, particularly in the bDH model. At the molecular level, selected inflammatory cytokines (IL-1α, IL-1β, Il-1ra, IL-6, and IL-17), chemokines (fractalkine, and in bDH only, RANTES/CCL5, IP-10/CXCL10), and adhesion molecules (sICAM-1) persisted or increased. Other inflammatory mediators declined. Chronic inflammation results from significant remodeling of the inflammatory response, it is not just a prolongation of the inflammatory dynamics initiated during the early stage.

The late increase in fractalkine (CX3CL1) is particularly relevant in the context of SCI. Fractalkine is constitutively expressed by CNS neurons, and signals primarily through CX3CR1 on microglia and macrophages. Following SCI, fractalkine/CX3CL1 is dynamically regulated, with an early decrease followed by a later increase (Donnelly et al., 2011; Kwiecien et al., 2020; present study, Fig.7). Notably, Kwiecien et al. (2020) suggested a potential regulatory role for this molecule during later stages after SCI, with its progressive late increase coinciding with astrogliosis and attenuation of several pro-inflammatory responses. In animals lacking fractalkine/CX3CR1, improved functional recovery after SCI and enhanced axonal sprouting and synaptogenesis are reported (Donnelly et al., 2011; Freria et al., 2017), indicating that the effects of CX3CL1/CX3CR1 signaling may depend on the cellular context and temporal evolution of SCI. Here, fractalkine peaked at 28 dpi (latest analyzed time point) in both lesion models, in accordance with previous observations (Hong et al., 2018; Kwiecien et al., 2020). This expression profile supports continued neuroimmune signaling as the lesion matures.

The persistence and remodeling of inflammatory activity may also be influenced by changes in the biomechanical properties of the lesion environment. The progressive formation of a glial scar and changes to the extracellular matrix composition are associated with tissue softening after SCI (Moeendarbary et al., 2017). Through mechanotransduction, tissue mechanics regulate immune cell activation, migration, phagocytosis, and cytokine production (Du et al., 2023). Thus, the distinct mechanical nature of contusion and bDH injuries may contribute to their respective early inflammatory signatures.

In conclusion, our work highlights the complexity of the inflammatory response following SCI, revealing similarities but also significant differences between a contusion and a bDH model, characterized by distinct initial mechanical impacts. Subsequent matrix remodeling, cavitation, and astroglial scar formation will progressively remodel the immune environment, potentially altering the inflammatory activity in the process. Finally, the results of this study may lay the foundation for therapeutic strategies aiming to modulate inflammation, reduce neurotoxicity, and promote tissue repair in SCI.

## ACKNOWLEDGMENTS

We thank the IBPS Imaging and animal facilities; Trung Hieu TRAN and Halley YUNG for contributing to histology experiments during their internships, Hélène Front for her invaluable support with the statistical analyses, and the SFAR (Société Française d’Anesthésie Réanimation) for providing KB’s internship fellowship.

## AUTHORS’ CONTRIBUTION

AG and CA, Investigation, Methodology, Visualization, Formal analysis, Data curation, Draft writing; KB, early conceptualization Investigation, Methodology, Visualization, Formal analysis; KK, Investigation, Methodology, Visualization; EKA and SS, Investigation, Formal analysis; vBY, Investigation, Methodology, Visualization, Formal analysis, Data curation, Writing, review, and Editing; FN, Conceptualization, Methodology, Investigation, Visualization, Formal analysis, Data curation, Validation, Supervision, Writing and Review, Project administration, Funding acquisition

## SUPPLEMENTARY FIGURES

**Supplementary figure 1:**
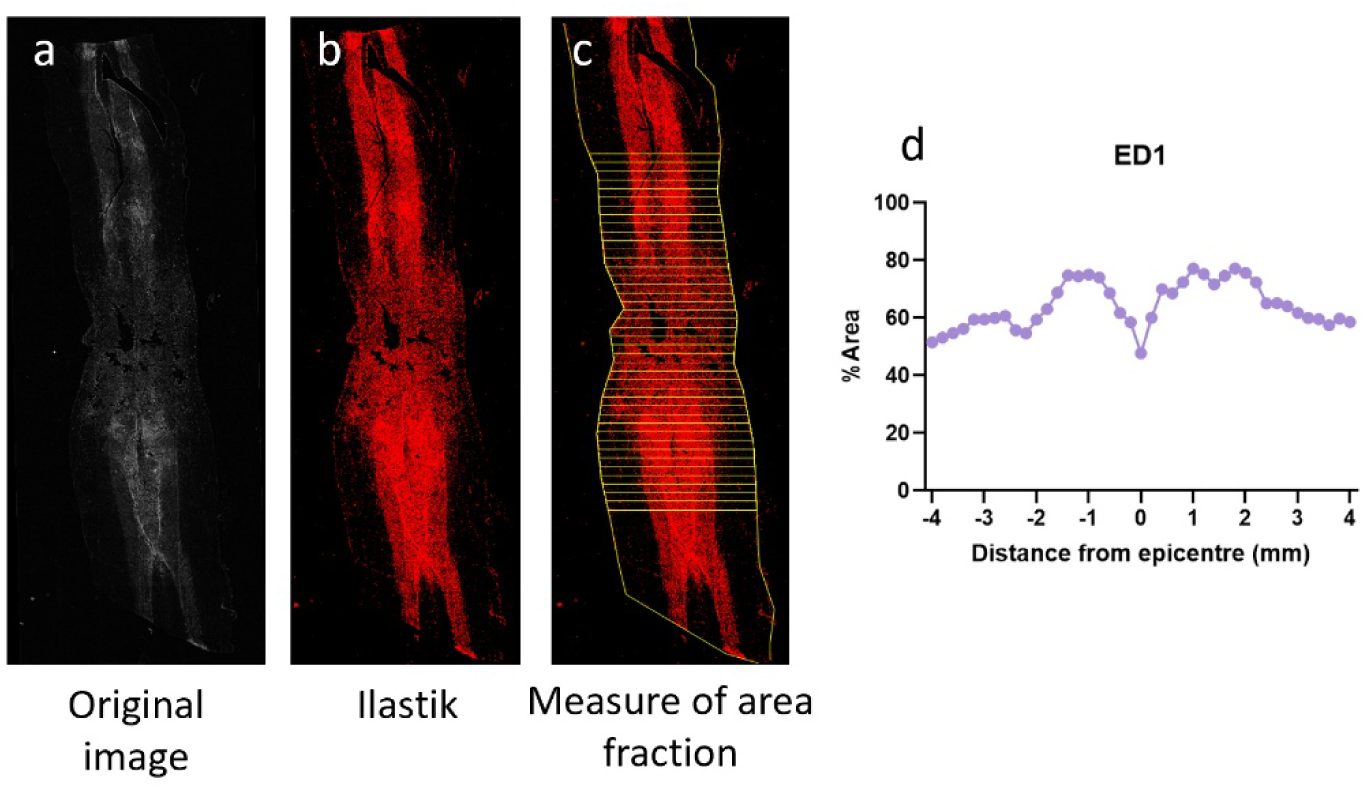
Method of quantitative analysis of ED1 and Iba-1 immunostaining through ROI segmentation Image Acquisition and Processing: Original microscope images were processed using a semi-automated pipeline composed of several steps: *Pre-processing and Contrast Enhancement*: Images were initially processed in ImageJ (Fiji) using the “Enhance Contrast” function, applying 0.35% saturation. Background subtraction was then performed using the “Subtract Background” function with a rolling ball radius of 50 pixels (a). *Segmentation via Supervised Pixel Classification:* Pre-processed images were segmented using Ilastik software (version 1.4.0) via interactive pixel classification, based on manual annotations. This process generated probability maps used for subsequent analyses (Berg et al., 2019). *Extraction of Channel of Interest:* Following Ilastik segmentation, the channel corresponding to the class of interest was extracted using a custom ImageJ macro script developed for this study (b). *Quantitative analysis based on Region of Interest (ROI) segmentation:* The final quantitative analysis was performed in ImageJ using a custom macro. For each image, the epicenter of the lesion was manually defined, and the left and right boundaries of the spinal cord section delimited. Based on these reference points, a series of 41 polygonal ROIs (each measuring 310 pixels, corresponding to 200 µm at the applied magnification; scale = 1.5501 pixels/µm) were automatically generated (c). These ROIs were used to quantify both the marked area and the area fraction (i.e., the percentage of the region invaded by Iba1 or ED1 immunofluorescence). In (d) is represented the result of quantification in (c).

**Supplementary figure 2:**
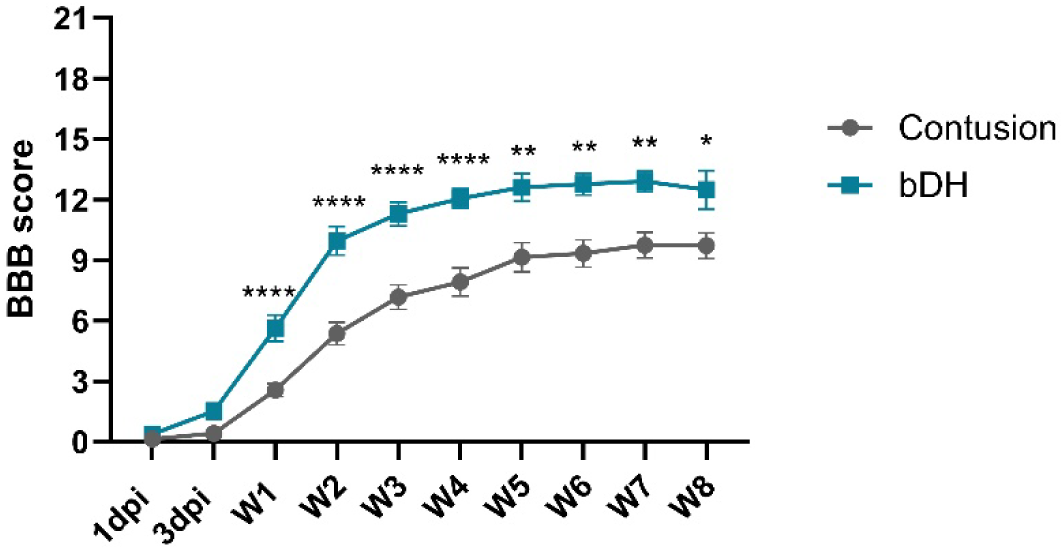
Spontaneous recovery of locomotor function was assessed one day and three days post-injury, then weekly for 8 weeks using the BBB locomotor rating scale. The BBB score of animals in the bDH group was higher as early as 3 dpi, becoming significant at one-week post-injury, compared with animals in the contusion group. This difference appears to be particularly important at 2, 3 and 4 weeks, and diminishes after this period, being no longer significant at 8 weeks post-injury. In both conditions, functional recovery appears to enter a stable state from 4 weeks post-injury (contusion, n = 71; bDH, n = 52). Stats: Two-way ANOVA, *p<0.05, **p<0.01, ***p<0.001, ****p<0.0001.

**Supplementary figure 3:**
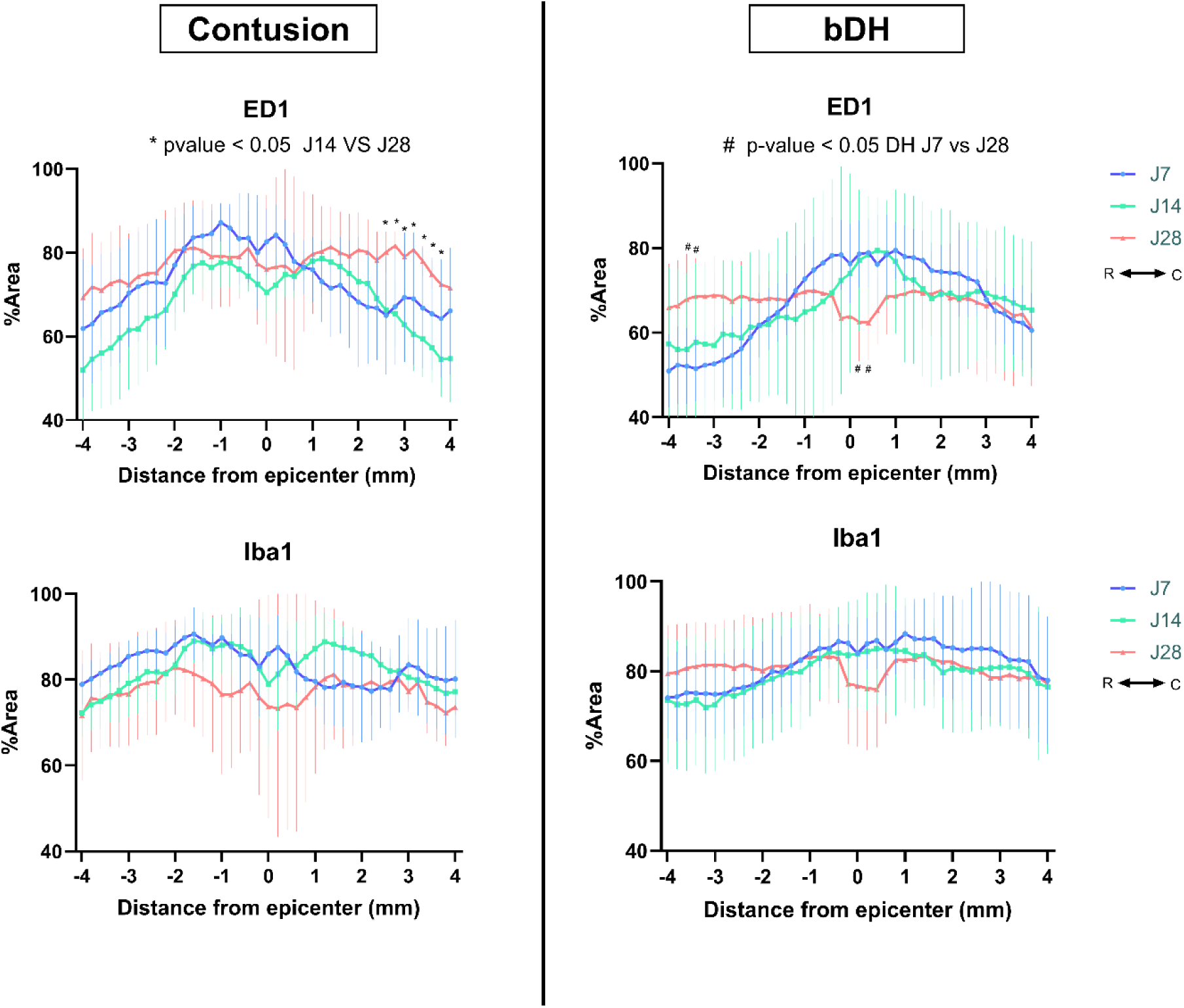
Temporal comparison of ED1 or Iba1 staining along the rostro-caudal axis in each injury model. ED1 and Iba1 immunostaining were quantitatively analyzed by segmenting the lesion region of interest (ROI) from the lesion epicenter (axis 0) to adjacent spinal cord segments, extending rostrally (0 to –4 mm) and caudally (0 to +4 mm), at 7, 14, and 28 dpi. For each marker and injury model, separate graphs illustrate the spatial distribution of staining along the rostro-caudal axis over time. In the bDH group, a significant increase in ED1 staining was observed rostrally at 28 dpi compared to 7 dpi, while a reduction in staining at the lesion epicenter was noted, likely reflecting cavity formation. In contrast, in the contusion group, ED1 staining was significantly higher caudally at 28 dpi compared to 14 dpi, suggesting a delayed and spatially distinct inflammatory response. For the contusion model, data were obtained from n=4 at 7 dpi, 5 at 14 dpi, and n=3 at 28 dpi. For the bDH model, n=5 at 7 dpi, 4 at 14 dpi, and n=5 at 28 dpi. Statistical analysis was based on two-way ANOVA followed by Tukey’s post hoc test; * or # indicate p < 0.05, and ** or ## indicate p < 0.01.

**Supplementary figure 4:**
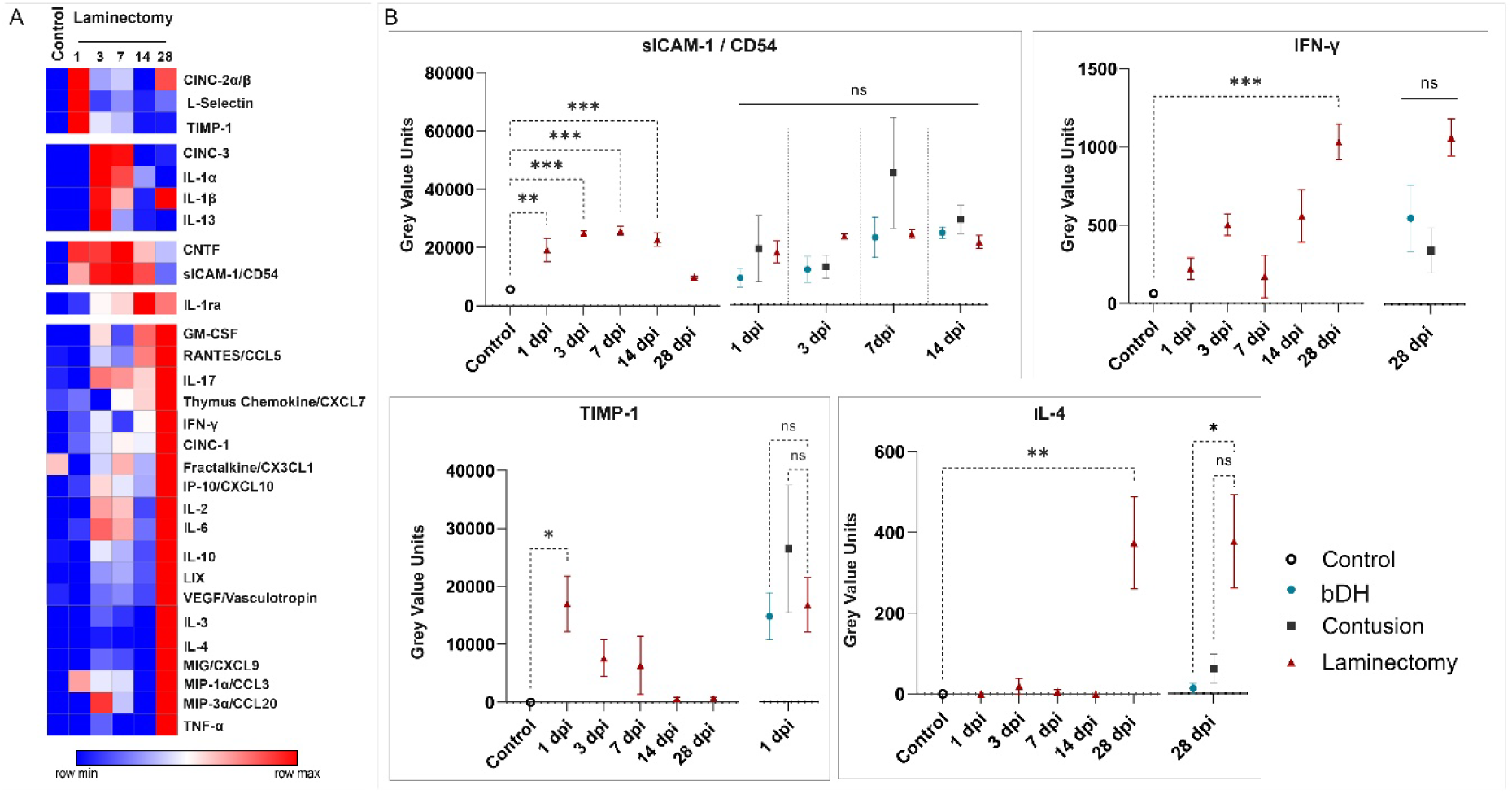
Comparative analysis of cytokine levels between Control and Laminectomy. **A.** Heatmap displaying temporal cytokine expression patterns in control and laminectomy groups. Data are derived from raw Gray Value Units and visualized using relative color coding, where blue indicates the minimum and red the maximum level. **B.** Temporal profiles of key cytokines differentially expressed between control (intact) and laminectomy groups (animals underwent laminectomy without SCI). The two-way ANOVA followed by Tukey’s post hoc analysis revealed significant differences for CD54 (1 dpi, 3 dpi, 7 dpi, 14 dpi), TIMP-1 (1 dpi), IL-4 (28 dpi), and IFN-g (28 dpi). Alongside, graphics show a comparison at the time points for which a significant difference was observed, between laminectomy, contusion, and bDH. Of note, only one significant difference was found for IL-4 at 28 dpi (p=0.0234). Error bars represent SEM. *p<0.05, **p<0.01, ***p<0.001, ****p<0.0001. Number of animals per condition: 3-4.

**Supplementary figure 5:**
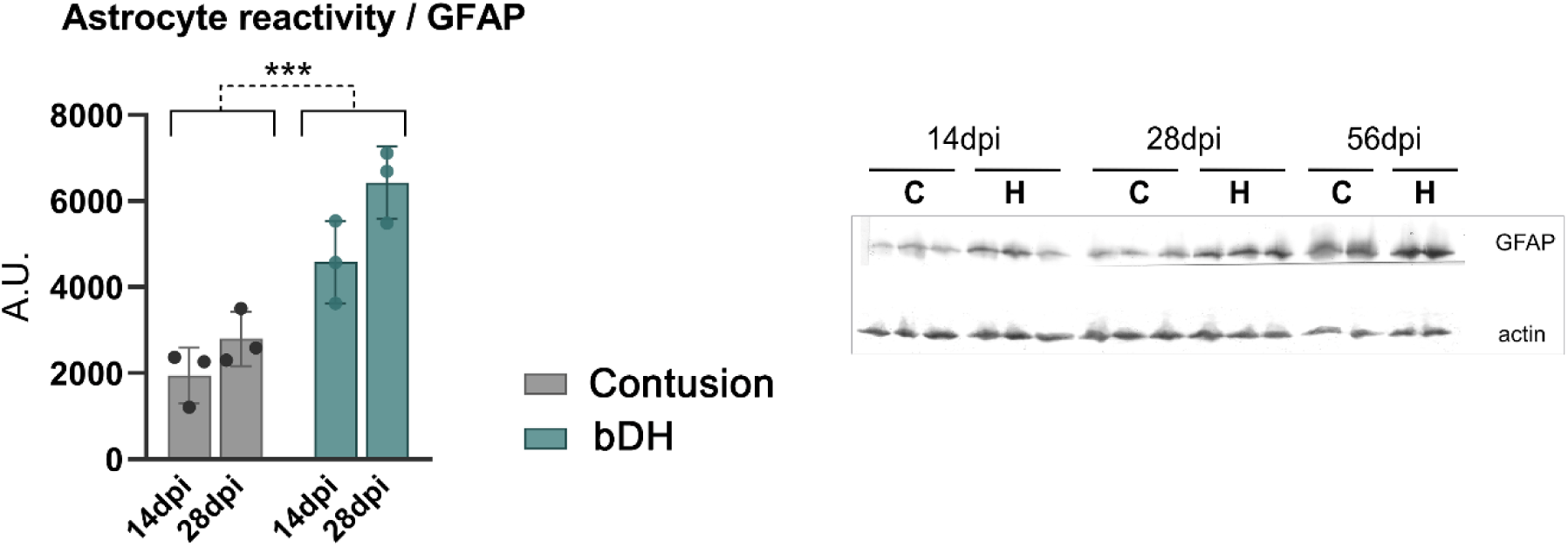
Western blot of spinal cord protein extracts labeled for reactive astrocyte marker GFAP (glial fibrillary acidic protein) at 14, 28 and 56 dpi (with actin staining used for normalization), and quantification of relative amounts of GFAP at 14 dpi and 28 dpi. GFAP expression is significantly higher in bDH compared to contusion group at 14 (**p<0.01) and 28 dpi (***p < 0.0001). Two-way ANOVA (n=3 per condition) revealed a main effect of time (p=0.0172) and of lesion type (p=0.0001), with no interaction of both.

